# The effect of network thresholding and weighting on structural brain networks in the UK Biobank

**DOI:** 10.1101/649418

**Authors:** Colin R. Buchanan, Mark E. Bastin, Stuart J. Ritchie, David C. Liewald, James Madole, Elliot M. Tucker-Drob, Ian J. Deary, Simon R. Cox

## Abstract

Whole-brain structural networks can be constructed using diffusion MRI and probabilistic tractography. However, measurement noise and the probabilistic nature of the tracking procedure result in an unknown proportion of spurious white matter connections. Faithful disentanglement of spurious and genuine connections is hindered by a lack of comprehensive anatomical information at the network-level. Therefore, network thresholding methods are widely used to remove ostensibly false connections, but it is not yet clear how different thresholding strategies affect basic network properties and their associations with meaningful demographic variables, such as age. In a sample of 3,153 generally healthy volunteers from the UK Biobank Imaging Study (aged 44—77 years), we constructed 85 × 85 node whole-brain structural networks and applied two principled network thresholding approaches (consistency and proportional thresholding). These were applied over a broad range of threshold levels across six alternative network weightings (streamline count, fractional anisotropy, mean diffusivity and three novel weightings from neurite orientation dispersion and density imaging) and for four common network measures (mean edge weight, characteristic path length, network efficiency and network clustering coefficient). We compared network measures against age associations and found that the most commonly-used level of proportional-thresholding from the literature (retaining 68.7% of all possible connections) yielded significantly weaker age-associations (0.070 ≤ |β| ≤ 0.406) than the consistency-based approach which retained only 30% of connections (0.140 ≤ |β| ≤ 0.409). However, we determined that the stringency of the threshold was a stronger determinant of the network-age association than the choice of threshold method and the two thresholding approaches identified a highly overlapping set of connections (ICC = 0.84) when matched at a plausible level of network sparsity (70%). Generally, more stringent thresholding resulted in more age-sensitive network measures in five of the six network weightings, except at the highest levels of sparsity (>90%), where crucial connections were then removed. At two commonly-used threshold levels, the age-associations of the connections that were discarded (mean β ≤ |0.068|) were significantly smaller in magnitude than the corresponding age-associations of the connections that were retained (mean β ≤ |0.219|, *p* < 0.001, uncorrected). Given histological evidence of widespread degeneration of structural brain connectivity with increasing age, these results indicate that stringent thresholding methods may be most accurate in identifying true white matter connections.

## Introduction

There has been a growing enthusiasm for work seeking to construct structural brain networks, or structural “connectomes” (Sporns et al., 2005), which map white matter connectivity between distal regions of the human brain. Structural connectomes can be estimated in vivo, at a macroscopic scale, using diffusion magnetic resonance imaging (dMRI) and whole-brain tractography (Sotiropoulos and Zalesky, 2017). This approach has been central for gauging how variation in the network organization of the brain relates to behaviour and health. However, the necessity of establishing a representative brain network for which within- and between-group comparisons can be conducted is a non-trivial task (de Reus and van den Heuvel, 2013). Due to the noisy and indirect measurement of water diffusion combined with the nature of probabilistic tractography, the resulting structural brain networks are known to contain many false-positive connections (Jbabdi and Johansen-Berg, 2011; Thomas et al., 2014; Yeh et al., 2018; Zalesky and Fornito, 2009). The unfiltered network generated by probabilistic tractography typically describes the brain as almost fully connected (Roberts et al., 2017). This is at odds with biological and post-mortem investigations of mammalian anatomical connectivity. Roberts et al. reported estimates of connection density varying from < 5% (Hagmann et al., 2008) to 13–36% for the entire brain and 32–52% for cortico-cortical connections in the mouse (Oh et al., 2014). Consequently, it may be inferred that many of the ‘connections’ identified in unfiltered networks are spurious.

Although previous research has been undertaken to map the major white matter pathways of the brain using dMRI and tractography (Mori et al., 2009), a fine-grained anatomical ‘ground-truth’, mapping the presence of every connection at the macroscale (typically thousands of white matter pathways involving millions of streamlines) has not yet been realised. A recent population-based atlas of white matter connectivity, constructed by manually labelling 40 major streamline clusters (Yeh et al., 2018), shows promise in validating connectivity but is limited to a small proportion of possible network connections.

In the absence of a comprehensive map of connectivity and to meet a demand for more principled network denoising approaches (de Reus and van den Heuvel, 2013; Maier-Hein et al., 2017; Van Wijk et al., 2010), researchers have introduced inferential methods to identify and discard potentially spurious connections. Some researchers have advocated using raw (unthresholded) matrices without removal of any connections on the basis that topological network properties are not significantly altered by the inclusion of weak connections (Civier et al., 2019). However, many network studies have employed thresholding strategies, such as, *absolute-thresholding* which applies a uniform threshold to retain only connections above a set weight (Hagmann et al., 2007), and *density-thresholding* which applies a (relative) threshold on the connection weights such that the weakest connections are removed to match the same number of connections across subjects (Rubinov and Sporns, 2010). Although the lowest weighted network connections (e.g., those involving fewest streamlines) are often false-positives, low weights do not necessarily correspond to implausible connections.

Consequently, more sophisticated thresholding approaches have been introduced which remove network connections using group-level statistics. *Proportional-thresholding* (consensus-thresholding) has been used to retain only the connections present in a set proportion of subjects (de Reus and van den Heuvel, 2013). Recent *consistency-thresholding* approaches have been introduced, which retain connections with weights that are consistent across subjects, on the assumption that connections with the highest inter-subject variability are spurious (Betzel et al., 2018; Roberts et al., 2017). Additionally, these approaches have involved schemes which promote white matter connections that are strong for their physical length to compensate for the bias in overestimating the number of short range connections (Betzel et al., 2018; Roberts et al., 2017). These studies also apply levels of network connection density based on evidence from tracing studies of the mammalian brain and thus may partly meet the need for more anatomically plausible measures of connectivity. However, a consensus on the thresholding method and range of threshold levels that might optimally reflect the underlying biological connectivity is currently lacking.

The lack of prior information on many of the thousands of possible connections in the brain has led researchers to evaluate the impact of different thresholding approaches using criterion validity – that is, the degree to which the network metrics are associated with external outcomes of interest. For example, a recent study examined four thresholding approaches in Huntington’s disease (138 participants), and found that detection of group-differences at the network level was highly dependent on the threshold level chosen (McColgan et al., 2018). However, the power limitation inherent in small samples restricts the fidelity with which the relative differences in validity across small changes in threshold level and network-weighting methods can be reliably detected.

We believe that age-associations, as a well-known correlate of white matter microstructure, are a strong candidate against which to determine the comparative criterion validity of network thresholding and weighting methods. Increasing age in adulthood is one of the most consistently replicated and widespread correlates of brain white matter macro- and microstructural outcomes (Bastin et al., 2010; Burzynska et al., 2010; Cox et al., 2016; Damoiseaux et al., 2009). Network analysis of alterations in brain organisation due to both normal and abnormal ageing have been applied to both structural and functional MRI (Alloza et al., 2018; Gong et al., 2009; Lo et al., 2010; Robinson et al., 2010; Zhao et al., 2015). Such studies have consistently demonstrated associations with brain-wide measures of connectivity and increasing age. Furthermore, histological studies of human and animal data have found evidence of the widespread degeneration of white matter (myelinated fibres) due to ageing and age-associated diseases (Salat, 2011).

In addition to the problem of spurious connections, there remains uncertainty about which network weighting best reflects the underlying biological connectivity. Various weights have been derived from dMRI structural networks, which reflect different notions of connection strength (Agosta et al., 2014; Collin et al., 2014; Hagmann et al., 2008; Robinson et al., 2010; Verstraete et al., 2011). The most common weightings used are interregional streamline counts/densities (Hagmann et al., 2008) and measures of water diffusion anisotropy (Robinson et al., 2010; Verstraete et al., 2011). Some network studies have introduced other dMRI weightings, such as mean diffusivity (MD), to characterise different aspects of white matter microstructure (Agosta et al., 2014; Collin et al., 2014). Neurite orientation dispersion and density imaging (NODDI; Zhang et al., 2012) provides a more sophisticated model of tissue microstructure than the conventional water diffusion tensor model (Cercignani and Bouyagoub, 2018). NODDI estimates neurite density (intra-cellular volume fraction; ICVF), extra-cellular water diffusion (isotropic volume fraction; ISOVF) and tract complexity/fanning (orientation dispersion; OD), biomarkers which can also be used as network weightings. Previous research has compared some conventional weightings (Buchanan et al., 2014; Dimitriadis et al., 2017; Qi et al., 2015), but it is not yet clear how thresholding affects differently-weighted networks and their relationships with external variables, such as age.

In the current study, using a large, single-scanner imaging sample (UK Biobank Imaging Study), we assessed the effect of two principled network thresholding approaches (proportional and consistency thresholding), both based on group-level statistics. We chose methods which operate at the group-level rather than the individual-level, because this matches the connection density across subjects and permits quantitative examination of individual differences. We exploited the large sample size to estimate the reproducibility of the both network thresholding methods in retaining the same connections across split-halves of the dataset. We also provided information with respect to the effect of thresholding across network weightings (streamline count, FA, MD, ICVF, ISOVF and OD) and four basic graph-theoretic measures (mean edge weight, characteristic path length, global network efficiency and network clustering coefficient).

We assessed the effect of thresholding using known age-associations in white matter (Cox et al., 2016). We first selected relevant threshold levels for each thresholding approach, based on the literature, for which to draw comparisons and to illustrate the practical implications for commonly-used thresholds. We then compared network-age associations over a range of threshold levels and assessed the individual age-associations of both retained and discarded connections. According to the principles underlying consistency and proportional thresholding, we hypothesised that: 1) thresholding would result in more age-sensitive network measures than unthresholded networks; 2) as the stringency of the threshold was increased, the magnitude of network-age associations would increase as more spurious connections were removed; and 3) the age-associations for discarded connections would be mainly null, based on the assumption that these were false-positive connections representing measurement noise.

## Methods

### Participants

The UK Biobank is a large-scale epidemiology study which recruited approximately 500,000 community-dwelling, generally healthy subjects aged 40—69 years from across Great Britain between 2006 and 2010. Participants provided comprehensive demographic, psychosocial and medical information during an initial visit to a UK Biobank assessment centre. Approximately 4 years after initial assessment a subset of participants underwent brain MRI (44—77 years of age) at the UK Biobank imaging centre in Cheadle, Manchester, UK. The initial release of dMRI data included 5,455 participants of whom 567 were excluded from the current study due to an incompatible dMRI acquisition used at an earlier scanning phase. A further 1,314 participants were removed by the UK Biobank following dMRI quality control procedures prior to release (as described in UK Biobank Brain Imaging Documentation). UK Biobank received ethical approval from the North West Multi-centre Research Ethics Committee (REC reference 11/NW/0382). All participants provided informed consent to participate. The current study was conducted under approved UK Biobank application number 10279.

### MRI acquisition and processing

Details of the MRI protocol and processing are freely available (Alfaro-Almagro et al., 2018; Miller et al., 2016a). All imaging data were acquired using a single Siemens Skyra 3T scanner. 3D T_1_-weighted volumes were acquired using a magnetization-prepared rapid gradient-echo sequence at 1 × 1 × 1 mm resolution with 208 × 256 × 256 field of view. The dMRI data were acquired using a spin-echo echo-planar imaging sequence (50 b = 1000 s/mm^2^, 50 b = 2000 s/mm^2^ and 10 b = 0 s/mm^2^) resulting in 100 distinct diffusion-encoding directions. The field of view was 104 × 104 mm with imaging matrix 52 × 52 and 72 slices with slice thickness of 2 mm resulting in 2 × 2 × 2 mm voxels.

Water diffusion parameters were estimated for FA, which measures the degree of anisotropic water molecule diffusion, and for MD, which measures the magnitude of diffusion. The parameters obtained from NODDI were: ICVF which measures neurite density; ISOVF which measures extracellular water diffusion; and OD which measures the degree of fanning or angular variation in neurite orientation (Zhang et al., 2012).

### Network construction

An automated connectivity mapping pipeline was used to construct white mater structural networks. This framework is described below with parameters informed by findings from a previous test-retest study using healthy volunteers (Buchanan et al., 2014).

Each T_1_-weighted image was segmented into 85 distinct neuroanatomical regions-of-interest (ROIs) using volumetric segmentation and cortical reconstruction (FreeSurfer v5.3.0). The Desikan-Killiany atlas was used to identify 34 cortical structures per hemisphere (Desikan et al., 2006). Subcortical segmentation was applied to obtain the brain stem and eight further grey matter structures per hemisphere: accumbens area, amygdala, caudate, hippocampus, pallidum, putamen, thalamus and ventral diencapahlon (Fischl et al., 2004, 2002). All cortical segmentations were visually quality checked for gross segmentation errors and 202 participants were removed.

A cross-modal nonlinear registration method was used to align ROIs from T_1_-weighted volume to diffusion space. Firstly, skull-stripping and brain extraction was applied to the FA volume of each participant (Smith, 2002). As an initial alignment, an affine transform with 12 degrees of freedom was used to align each brain-extracted FA volume to the corresponding FreeSurfer extracted T_1_-weighted brain using a mutual information cost function (FLIRT; Jenkinson and Smith, 2001). Local alignment was then refined using a nonlinear deformation method (FNIRT; Andersson et al., 2007). FreeSurfer segmentations were then aligned to diffusion space using nearest neighbour interpolation. For each participant, a binary mask used to constrain tractography, was formed in diffusion space from all grey and white matter voxels.

Whole-brain tractography was performed using an established probabilistic algorithm and a two-fibre model (BEDPOSTX/ProbtrackX; Behrens et al., 2007, 2003). Probability density functions, which describe the uncertainty in the principal directions of diffusion, were computed with a two-fibre model per voxel (Behrens et al., 2007). Streamlines were then constructed by sampling from these distributions during tracking using 100 Markov Chain Monte Carlo iterations with a fixed step size of 0.5 mm between successive points. Tractography was initiated from all white matter voxels and streamlines were constructed in two collinear directions until terminated by the following stopping criteria: 1) exceeding a curvature threshold of 70 degrees; 2) entering a voxel with FA below 0.1; 3) entering an extra-cerebral voxel; 4) exceeding 200 mm in length; and 5) exceeding a *distance ratio metric* of 10. This tracking criteria was set to minimize the amount of anatomically implausible streamlines. The distance ratio metric (Bullitt et al., 2003), excludes implausibly tortuous streamlines, for which a streamline with a length ten times longer than the distance between end points was considered invalid.

Networks were constructed by identifying connections between all ROI pairs. The endpoint of a streamline was recorded as the first ROI encountered (if any) when tracking from the seed location. Successful connections were recorded in an 85 × 85 connectivity matrix. A network weighting based on absolute streamline count (SC) was computed, *a_ij* = *count(i, j)*, which is the count of all streamlines identified between nodes *i* and *j*. In order to apply the streamline length correction (Roberts et al., 2017), a length matrix, *d_ij* was computed for each participant recording the mean length along all interconnecting streamlines between node *i* and *j*. A group-wide matrix of mean streamline lengths, *l*, was then constructed by taking the element-wise mean across the set of length matrices. In addition to the streamline count, five further network weightings were computed for FA, MD, ICVF, ISOVF and OD. For each weighting, a connectivity matrix was computed with element, *a_ij*, recording the mean value of the diffusion parameter in voxels identified along all interconnecting streamlines between nodes *i* and *j*. As tractography cannot distinguish between afferent and efferent connections, all matrices were made symmetric. Self-connections (diagonal elements) were removed and set to zero for all matrices.

### Network thresholding and network measures

We applied both proportional and consistency thresholding using SC-weighted networks, which reflect the likelihood of connection obtained from probabilistic tractography. Proportional-thresholding was applied to the set of SC matrices by only retaining the network connections which occurred (i.e., were nonzero) in a given proportion of subjects. Consistency-thresholding was applied by first correcting each SC matrix by the length matrix, *a_ij*/*l_ij*, in order to correct for the bias in identifying short-range connections (Roberts et al., 2017). A threshold on the coefficient of variation (CoV) of the length normalised weights was then applied to retain a set of connections across subjects. As the threshold criteria for the proportional and consistency approaches were not directly comparable, we measured both against network sparsity (the proportion of zero-weighted elements out of the total number of possible elements in a connectivity matrix).

For each approach, we selected a relevant threshold level for which to draw comparisons. For proportional-thresholding this level was set as the proportion of connections that were present in at least 50% of subjects, based on the median value obtained from a literature search of network studies published in the last two years (Supplementary Table 1). The consistency-thresholding level was set to retain 30% of connections, based on estimates from human and animal in vivo and in vitro data (Roberts et al., 2017). In addition to these levels, networks were computed over 100 equally spaced threshold levels from 0 to 100% network sparsity. For each threshold level, the same threshold mask was applied to each of the five diffusion weighted networks resulting in identical network sparsity across weightings.

For each network weighting (SC, FA, MD, ICVF, ISOVF and OD) and threshold level, per subject, we computed the *mean edge weight* (mean of all network connections including any zero-weighted elements which survived group-wide thresholding). In addition, three global graph-theoretic metrics were computed (Rubinov and Sporns, 2010): *characteristic path length* (a measure of network integration), *global network efficiency*, and *network clustering coefficient* (reflecting the interconnectedness of each node’s neighbours).

### Statistical analysis

We first computed the mean connectivity matrices and reported descriptive statistics for unthresholded networks across the six network weightings. Following network thresholding, we computed correlations between each of the four network measures for six weightings and for three threshold levels. Similarly, correlations between the six weightings, in terms of mean edge weight, were computed over the same threshold levels.

We computed the group-level statistics used by both thresholding methods and provided details on the proportion of subjects in which connections occurred and the number of streamlines involved. We measured the reproducibility of both thresholding approaches by measuring how consistently each retained the same network connections using different subsets of the full cohort. Split-half agreement was computed by randomly splitting the full dataset into halves (N = 1,577 and N = 1,576), computing two independent thresholds in each half and then using the intraclass correlation coefficient (ICC) to compute the agreement (presence of connections) in matrix elements (85 × 85 binary matrices) obtained from the two thresholds. We computed ICC(3,1), which applies a two-way mixed model with single measures and consistency of agreement (Shrout and Fleiss, 1979). At each of the 100 threshold levels, this split-half agreement procedure was repeated by resampling 1,000 times to compute mean values and confidence intervals.

We used age-associations as a test bed to compare the criterion validity of proportional and consistency thresholding. Initially, we compared the two methods at the pre-defined threshold levels alongside unthresholded matrices: connections present in 50% of participants (PT50); and consistency-thresholding at 30% (CT30). We therefore have three thresholding configurations for comparison, termed Raw, PT50, and CT30. We recognise that the these configurations have different network sparsities (0.313 for PT50; and 0.700 for CT30), and thus also provide two additional comparators matched for sparsity (PT at 0.700 sparsity, and CT at 0.313 sparsity). This allows us to comment on the relative contributions of thresholding method (proportional/consistency) or general stringency of the threshold (i.e., network sparsity) to observed network-age associations. For the set threshold levels (PT50 and CT30 and their sparsity-matched corollaries), we tested for the difference between a pair of age-associations based on dependent groups (Williams, 1959). We then provided a more in-depth analysis by testing both proportional and consistency thresholding over a range of threshold levels.

To assess the efficacy of thresholding in a sample with known age-associations we extracted the network connections that were retained and discarded, to test the hypothesis that discarded connections would be spurious and would therefore exhibit mainly null age-associations, and that the increased signal-to-noise in retained connections would result in stronger age-associations. Unpaired two-sample *t*-tests were used to test the difference in age-associations between the set of connections that were retained and the corresponding connections that were discarded. We compared properties between these two classes in terms of age-associations (standardised betas) of the mean edge weights. In order to visualise the regions involved for both retained and discarded connections computed with CT30, anatomical circle plots were constructed (Irimia et al., 2012), which grouped related neuroanatomical nodes and plotted connections by strength of age-association.

Throughout, for each of the six weightings and four network measures, multiple regression was used to model the associations between network measures with respect to age, age^2^, sex and age × sex. Owing to the large number of comparisons a threshold of *p* < 0.001 (uncorrected) was used to denote significant effects in each model. Uncorrected p-values were reported because our intention was to use the known age-association in white matter as a comparator (given the absence of ground-truth data) between different weightings and thresholds. The above associations (standardised betas), standard error and adjusted R^2^ were computed for each weighting over all 100 threshold levels.

We restricted our main analyses, throughout, to streamline counts that were uncorrected for grey or white matter volumes. At a constant resolution, network methods may identify more inter-regional streamlines in large brains than in small brains, but some have suggested that volume correction of streamline counts may overcompensate for volume-driven effects on these streamline weightings (Van Den Heuvel and Sporns, 2011). In a supplementary analysis, we therefore investigated differences in the criterion validity (for both age and sex, given the well-replicated sex differences in brain size; Ritchie et al., 2018) of mean edge weight when applying four variants of SC-weighting: uncorrected streamline count; network-wise correction by the number of seed points per subject (count of white matter voxels); streamline density with edge-wise correction by the count of voxels per ROI (Hagmann et al., 2008); and streamline density with edge-wise correction by node surface area at the white matter interface (the count of voxels which directly neighbour a white matter voxel).

## Results

### Network characteristics

3,153 participants (44.6–77.1 years of age, 1,496 male) remained after participants were excluded at quality checking or due to failure in processing. On average, 6.01 million streamlines were seeded per subject of which 1.49 million (24.9%) were found to successfully connect between nodes following the tracking procedure and removal of self-connections. The mean connectivity matrices and corresponding histograms of edge weights computed for each network weighting (SC, FA, MD, ICVF, ISOVF and OD) are shown in Figure 1. In each case, the networks were produced from the same set of streamlines. Before any thresholding, the mean value of network sparsity (the proportion of zero-weighted elements in a connectivity matrix) across subjects was 0.316 (SD = 0.032) meaning that probabilistic tractography generated brain networks with a connection density of ∼68% per subject. The mean connectivity matrix computed across all subjects (Figure 1) was almost fully connected with a sparsity of 0.002, although 69.8% of all connections in this network involved fewer than 50 streamlines per subject on average. We observed from the histograms of edge weights pooled across all subjects (Figure 1) that the distribution of SC weights approximately followed a power law and involved many low weighted connections but very few high weighted connections, whereas the five dMRI-based weightings followed approximately normal distributions.

**Figure 1.**
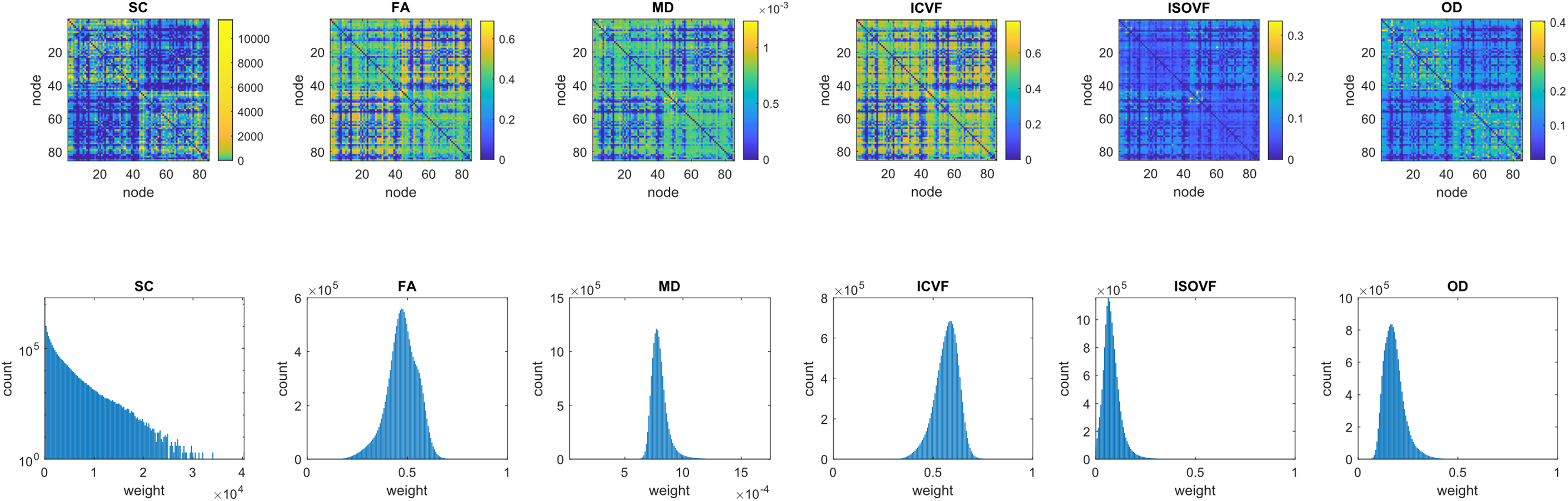
Top: 85 × 85 mean connectivity matrices (unthresholded) of inter-region connection weights averaged across all participants (N = 3,153) for six network weightings and generated from the same set of streamlines. In each case, the two large rectangular patterns on the diagonal correspond to the left and right hemispheres. Bottom: the corresponding histograms of nonzero edge weights pooled across all participants for each weighting (SC is log-scaled). SC = streamline count, FA = fraction anisotropy, MD = mean diffusivity, *ICVF = Intracellular volume fraction, ISOVF = Isotropic volume fraction, OD = orientation dispersion.*

The absolute values of graph-theoretic metrics are known to be dependent on the network methods used (Qi et al., 2015). Whilst the absolute values of our global network measures (mean edge weight, characteristic path length, network efficiency and network clustering coefficient) are not key to our analyses, we found that the value of each measure varied considerably across weightings and sparsities (Supplementary Table 2). Generally, as network sparsity was increased, both mean edge weight and characteristic path length also increased, whereas both network efficiency and network clustering coefficient decreased. We found that our four global network measures were highly correlated with each other when assessed for each of the six network weightings and for three threshold levels (Raw, PT50 and CT30), with all correlations between these network measures at *r* ≥ |0.60| for SC-weighted networks and *r* ≥ |0.74| for all other weightings. Similarly, the mean edge weights of the six network weightings were found to be highly correlated in several cases (Supplementary Figure 1). For example, at the most stringent level of sparsity tested (CT30), the strongest correlations were between FA and ICVF (*r* = 0.84); MD and ICVF (*r* = -0.83); FA and MD (*r* = -0.73); MD and OD (*r* = -0.46); and MD and ISOVF (*r* = 0.44).

### Comparison of thresholding methods

To illustrate the group-level statistics used by the proportional and consistency thresholding approaches, we calculated the proportion of subjects in which each connection existed (Figure 2A) and the inter-subject variability (CoV) in streamline count following length correction (Figure 2B). The elements of the proportional and CoV matrices were significantly correlated at *r* = -0.61. The relationship between the threshold on the proportion of subjects and the resulting network sparsity is nonlinear (Figure 2C). The relationship between the threshold on the inter-subject variability of edge weights and network sparsity (Figure 2D) shows that there are few connections with high inter-subject variability (CoV > 10) and most connections exhibit low inter-subject variability (e.g., CoV < 1.25 for the top 30% most consistent connections). Proportional-thresholding could not be applied at the highest levels of network sparsity because above 81% sparsity no further connections could be removed by this criterion, i.e., 19% of all possible network connections were present in every subject.

**Figure 2.**
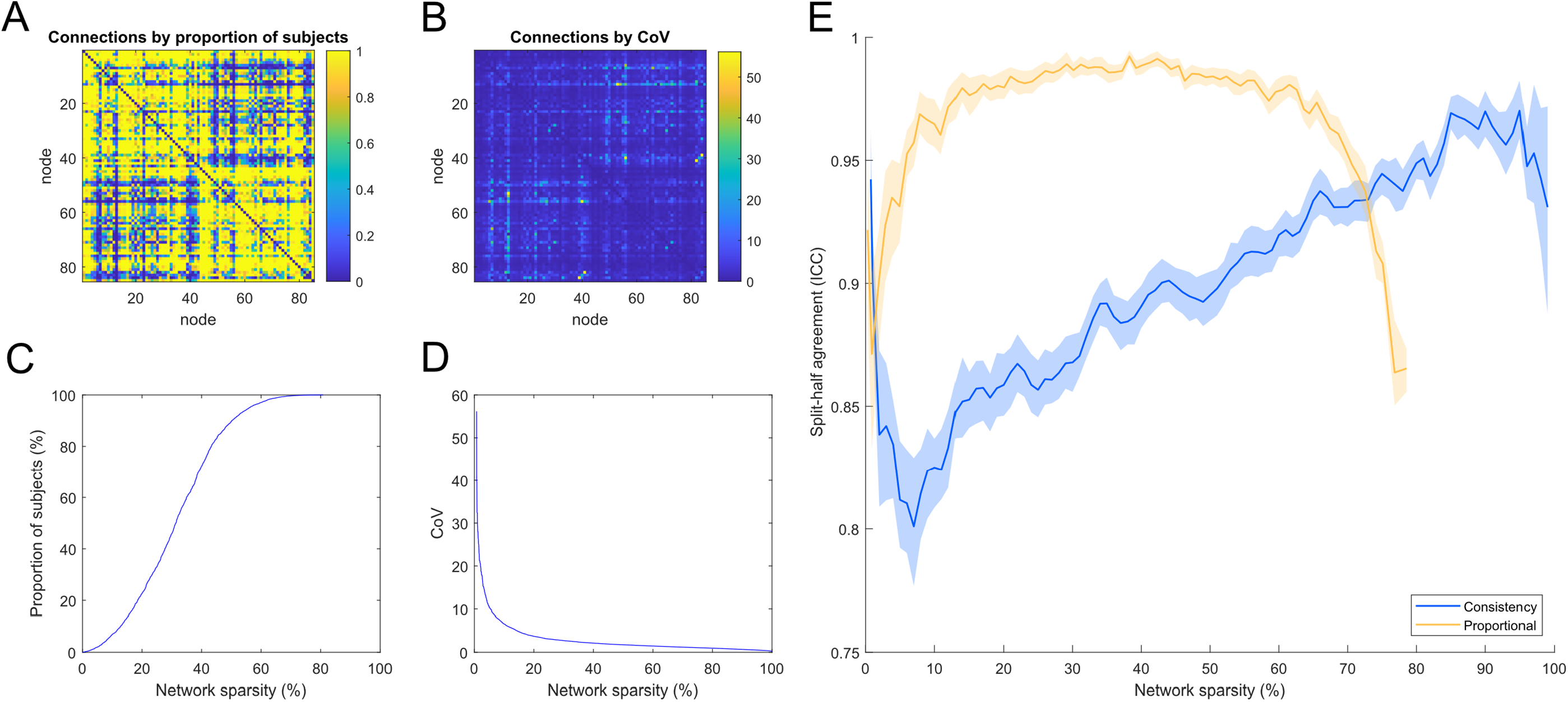
A) 85 × 85 matrix showing the proportion of subjects for which each network connection was present; B) 85 × 85 matrix showing the coefficient of variation (CoV) in network weights following streamline count length correction used by consistency-thresholding; C) The relationship between the threshold on the proportion of subjects and network sparsity; D) The relationship between the threshold on CoV of length-corrected weights and network sparsity; E) Split-half agreement of thresholded connections computed by: randomly splitting the dataset into halves, applying two independent thresholds and using ICC to measure the agreement (presence of connections) identified from the two thresholds (95% CIs computed over 1000 resampling iterations and plotted against network sparsity).

Whereas CT30 retained 30.0% of 3,570 possible network connections, the threshold level for PT50 was more relaxed and retained 68.7% of all connections. Notably, the network connections removed by either thresholding method involved relatively few streamlines per subject. For instance, CT30 removed 2,493 (70.0%) of the network connections but this only discarded 75 thousand streamlines per subject, and the majority of streamlines (approximately 1.42 million) were retained. Similarly, PT50 removed 1,119 (31.3%) of the network connections but this only discarded ∼650 streamlines per subject (i.e., discarded connections had a streamline count of zero in many subjects), and approximately 1.49 million streamlines were retained. Therefore, most connections removed by either thresholding method mainly comprised very few streamlines. This is consistent with the hypothesis that such connections might be regarded as spurious.

For both thresholding approaches, measuring how consistently the same network connections were identified in separate halves of the sample, after random splitting into halves (N = 1,577 and N = 1,576) and computing two independent thresholds, resulted in high agreement (mean ICCs > 0.81; Figure 2E). The agreement for PT50 (mean ICC = 0.99) was greater than the agreement for CT30 (mean ICC = 0.93). When examined across the full range of sparsities, proportional-thresholding was highly consistent (mean ICCs > 0. 97) over 20 to 60% sparsity but agreement declined above 60% sparsity, presumably as core white matter connections were pruned from one sample but not the other. Consistency-thresholding showed a broadly linear increase in ICC scores as connections were removed with the highest mean ICC of 0.97 obtained with 5% of connections remaining. Although the proportional approach obtained higher agreement than the consistency approach over most levels, we observed that the crossover in their performance occurred at around 70% sparsity (coinciding with the 30% of connections threshold level proposed by Roberts et al., 2017). Similarly, when both thresholding methods were compared across the full sample when measured at a matched sparsity of 70% (CT30 and PT at 0.7 sparsity), the agreement in the network connections identified by the two methods was high (ICC = 0.84 or 948/1,190 matching connections).

### Network-age associations by thresholding method

We examined the age-associations of global network measures obtained for PT50 (corresponding to a network sparsity of 31.3%) and CT30 (sparsity of 70.0%), alongside results from the set of Raw matrices. In order to parse apart the contributions of network sparsity to any observed differences between thresholding methods, we also provided age-associations for two other threshold levels, for which network sparsity was matched between threshold methods. The linear component for age-associations between four metrics (mean edge weight, characteristic path length, network efficiency and network clustering coefficient) from each of the six weightings (SC, FA, MD, ICVF, ISOVF, OD) across these thresholding methods are shown in Table 1 and Figure 3. Full regression coefficients (age, age^2^, sex and sex × age) are presented in Supplementary Tables 3-5.

**Figure 3.**
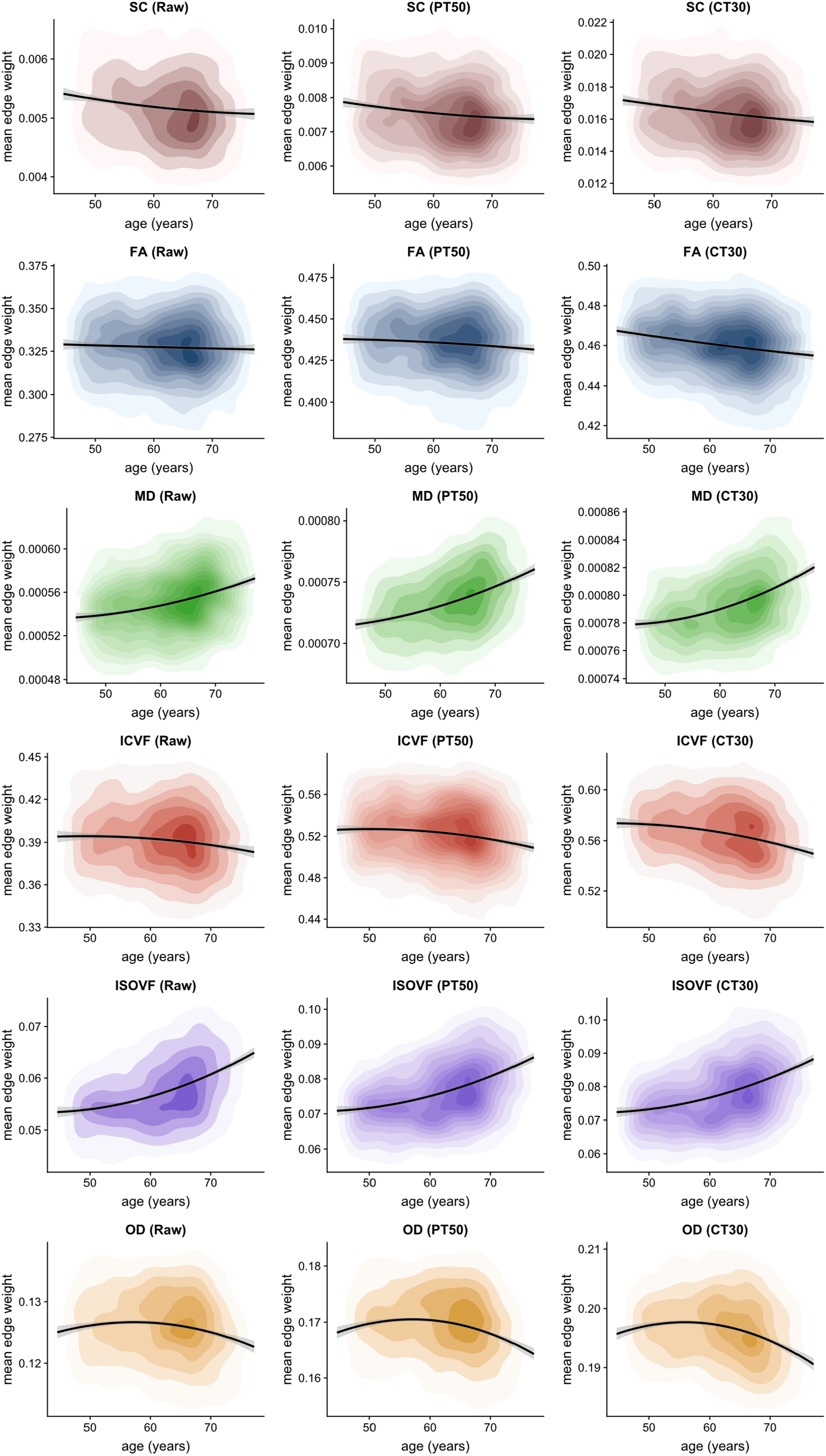
Density plots of mean edge weight for six network weightings under three alternative thresholding approaches: unthresholded (Raw), proportional-thresholding at 50% of subjects (PT50) and consistency-thresholding at 30% (CT30), with quadratic fit and 95% CIs.

**Table 1.** Comparison of age-associations (standardised regression coefficients and uncorrected *p*-values) for four network metrics measured across six network weightings and five thresholding approaches. The five thresholding approaches are: Raw (unthresholded); proportional at 50% (PT50), consistency at 68.6% (matched on network sparsity to PT50 for comparison), proportional at 99.6% (matched on sparsity with CT30) and consistency at 30% (CT30). The reported regression coefficients denote the linear component for age and full regression coefficients (age, age^2^, sex and sex × age) are presented in Supplementary Tables 3-5. Emboldening indicates significance (*p* < 0.001, uncorrected). Tests for the difference in the magnitude of age-associations were conducted for Raw vs. PT50 and PT50 vs. CT30; all were significantly different apart from _a_ and _b_, which denote nonsignificantly different pairs of magnitudes (*p* < 0.001, uncorrected). SC = streamline count, FA = fraction anisotropy, MD = mean diffusivity, ICVF = Intracellular volume fraction, ISOVF = Isotropic volume fraction, OD = orientation dispersion.

| Network metric | Weighting | Raw | PT50:<br>Proportional<br>(50%) | Consistency<br>(68.6%) | Proportional<br>(99.6%) | CT30:<br>Consistency<br>(30%) |
| --- | --- | --- | --- | --- | --- | --- |
|  |  | sparsity = 0.002 | sparsity = 0.313 | sparsity = 0.313 | sparsity = 0.701 | sparsity = 0.700 |
| Mean edge weight | SC | **-0.141 (<0.001)** | **-0.141 (<0.001)** | **-0.142 (<0.001)** | **-0.154 (<0.001)** | **-0.172 (<0.001)** |
|  | FA | -0.033 (0.079) | **-0.070 (<0.001)** | **-0.069 (<0.001)** | **-0.187 (<0.001)** | **-0.179 (<0.001)** |
|  | MD | **0.285 (<0.001)** | **0.346 (<0.001)<sup>b</sup>** | **0.340 (<0.001)** | **0.381 (<0.001)** | **0.378 (<0.001)<sup>b</sup>** |
|  | ICVF | **-0.097 (<0.001)** | **-0.134 (<0.001)** | **-0.133 (<0.001)** | **-0.212 (<0.001)** | **-0.206 (<0.001)** |
|  | ISOVF | **0.362 (<0.001)<sup>a</sup>** | **0.367 (<0.001)<sup>a</sup>** | **0.367 (<0.001)** | **0.351 (<0.001)** | **0.353 (<0.001)** |
|  | OD | **-0.101 (<0.001)** | **-0.146 (<0.001)** | **-0.146 (<0.001)** | **-0.198 (<0.001)** | **-0.198 (<0.001)** |
| Characteristic path length | SC | **0.160 (<0.001)<sup>a</sup>** | **0.160 (<0.001)<sup>ab</sup>** | **0.160 (<0.001)** | **0.129 (<0.001)** | **0.160 (<0.001)<sup>b</sup>** |
|  | FA | **0.089 (<0.001)** | **0.109 (<0.001)** | **0.114 (<0.001)** | **0.152 (<0.001)** | **0.140 (<0.001)** |
|  | MD | **-0.332 (<0.001)** | **-0.358 (<0.001)<sup>b</sup>** | **-0.361 (<0.001)** | **-0.378 (<0.001)** | **-0.364 (<0.001)<sup>b</sup>** |
|  | ICVF | **0.149 (<0.001)** | **0.167 (<0.001)** | **0.167 (<0.001)** | **0.185 (<0.001)** | **0.184 (<0.001)** |
|  | ISOVF | **-0.339 (<0.001)<sup>a</sup>** | **-0.341 (<0.001)<sup>a</sup>** | **-0.328 (<0.001)** | **-0.269 (<0.001)** | **-0.290 (<0.001)** |
|  | OD | **0.217 (<0.001)** | **0.229 (<0.001)** | **0.225 (<0.001)<sup>b</sup>** | **0.216 (<0.001)** | **0.219 (<0.001)<sup>b</sup>** |
| Network efficiency | SC | **-0.186 (<0.001)<sup>a</sup>** | **-0.186 (<0.001)<sup>a</sup>** | **-0.186 (<0.001)** | **-0.186 (<0.001)** | **-0.192 (<0.001)** |
|  | FA | **-0.095 (<0.001)** | **-0.117 (<0.001)** | **-0.118 (<0.001)** | **-0.168 (<0.001)** | **-0.158 (<0.001)** |
|  | MD | **0.385 (<0.001)** | **0.405 (<0.001)<sup>b</sup>** | **0.407 (<0.001)** | **0.418 (<0.001)** | **0.409 (<0.001)<sup>b</sup>** |
|  | ICVF | **-0.160 (<0.001)** | **-0.176 (<0.001)** | **-0.175 (<0.001)** | **-0.201 (<0.001)** | **-0.197 (<0.001)** |
|  | ISOVF | **0.380 (<0.001)** | **0.389 (<0.001)<sup>b</sup>** | **0.392 (<0.001)** | **0.396 (<0.001)** | **0.401 (<0.001)<sup>b</sup>** |
|  | OD | **-0.217 (<0.001)<sup>a</sup>** | **-0.225 (<0.001)<sup>ab</sup>** | **-0.223 (<0.001)** | **-0.215 (<0.001)** | **-0.215 (<0.001)<sup>b</sup>** |
| Network clustering coefficient | SC | **-0.098 (<0.001)<sup>a</sup>** | **-0.087 (<0.001)<sup>a</sup>** | **-0.096 (<0.001)** | **-0.143 (<0.001)** | **-0.179 (<0.001)** |
|  | FA | -0.056 (0.002) | **-0.107 (<0.001)** | **-0.112 (<0.001)** | **-0.198 (<0.001)** | **-0.192 (<0.001)** |
|  | MD | **0.409 (<0.001)<sup>a</sup>** | **0.406 (<0.001)<sup>a</sup>** | **0.399 (<0.001)** | **0.374 (<0.001)** | **0.368 (<0.001)** |
|  | ICVF | **-0.127 (<0.001)** | **-0.162 (<0.001)** | **-0.165 (<0.001)** | **-0.214 (<0.001)** | **-0.209 (<0.001)** |
|  | ISOVF | **0.345 (<0.001)** | **0.338 (<0.001)** | **0.336 (<0.001)** | **0.313 (<0.001)** | **0.311 (<0.001)** |
|  | OD | **-0.142 (<0.001)** | **-0.178 (<0.001)<sup>b</sup>** | **-0.186 (<0.001)** | **-0.178 (<0.001)** | **-0.178 (<0.001)<sup>b</sup>** |

First, we compared age-associations for Raw vs. PT50 vs. CT30 (columns 1, 2 and 5 in Table 1). The overarching pattern of results indicated that age-association magnitudes were Raw < PT50 < CT30, with effect sizes (magnitude of standardised regression coefficients) between 0.033 and 0.409 for Raw, between 0.070 and 0.406 for PT50 and between 0.140 and 0.409 for CT30. The differences between pairs of age-associations over these levels were significant in the majority of cases (*p* < 0.001, uncorrected; the 15 nonsignificant instances are indicated in Table 1). Over these three levels, FA-weighted networks showed the most pronounced increase in effect size as sparsity was increased (0.033 < 0.070 < 0.179). However, when the thresholding methods were matched by sparsity (columns 3 and 4 in Table 1), it became apparent that network sparsity rather than the specific thresholding approach was the main driver of differences in age-associations with the various network metrics. Excluding SC and ISOVF, generally the magnitude of the age-association increased as the stringency of the threshold was increased (and more uninformative network connections were removed). In most cases, metrics derived from Raw matrices yielded the weakest age-associations (0.033 ≤ |β| ≤ 0.409). However, there was little evidence that consistency-thresholding yielded stronger age-associations than the proportional method when sparsity was matched.

Although various interregional streamline count weightings have previously been used, we have limited our analysis above to uncorrected streamlines counts. In a supplementary analysis, we also computed three other streamline network weightings and computed age and sex associations using mean edge weight (Supplementary Table 6). As expected, increasing age was associated with fewer streamlines for uncorrected SC (−0.172 ≤ β ≤ -0.141, *p* < 0.001), but also with being male (0.379 ≤ β ≤ 0.384, *p* < 0.001), at all thresholds tested. When SC was corrected for the number of white matter seed points per subject, there was no significant age-association (−0.005 ≤ β ≤ 0.042, all nonsignificant), and females had greater streamline weights (−0.173 ≤ β ≤ -0.159, *p* < 0.001). When correcting streamline count for grey matter node volume or surface area, age-associations were flipped, denoting higher weights with increasing age (0.067 ≤ β ≤ 0.173, *p* < 0.001), and yielding null sex differences except for Raw and PT50 thresholded networks using surface area correction (β = 0.082, *p* < 0.001 in both cases), indicating males had modestly higher weights. No significant sex effects were found with the dMRI weightings when measuring mean edge weight (Supplementary Tables 3-5).

### Network-age associations by thresholding level

Age-associations for retained versus discarded connections were also compared across the entire sparsity range from 0 to 100% (Figure 4). When visualised across all threshold levels, the profile of age-associations across thresholds levels (both retained and discarded connections) were very similar between proportional and consistency-thresholding. However, age-associations for the proportional method could not be computed at the highest levels of sparsity because of the limits of the thresholding criterion. Generally, more stringent thresholding (increasing sparsity) resulted in a greater magnitude of age-association for retained connections in SC, FA, MD, ICVF and OD, except at the highest levels of sparsity (>90%) where crucial connections were then removed (though for ISOVF, increasing the threshold level resulted in a slight decrease in the magnitude of the age-association for retained connections). Whereas the range for the strongest age-association in retained connections was approximately 50–95% sparsity (across weightings), age-associations for the discarded connections were closest to null below 50% sparsity. Crucially, it was observed that at nearly every threshold level the magnitude of the age-associations for retained connections was greater than the corresponding age-association for the discarded connections (except for SC). The discarded connections showed null profiles for FA, ICVF and OD (but not for SC, MD and ISOVF) across most of the sparsity range. Although the magnitudes of age-associations were larger for retained versus discarded connections, both were significant, and in the same direction, across most levels of network sparsity; for ISOVF the age-association became broadly equivalent for both retained and discarded connections above 70% sparsity.

**Figure 4.**
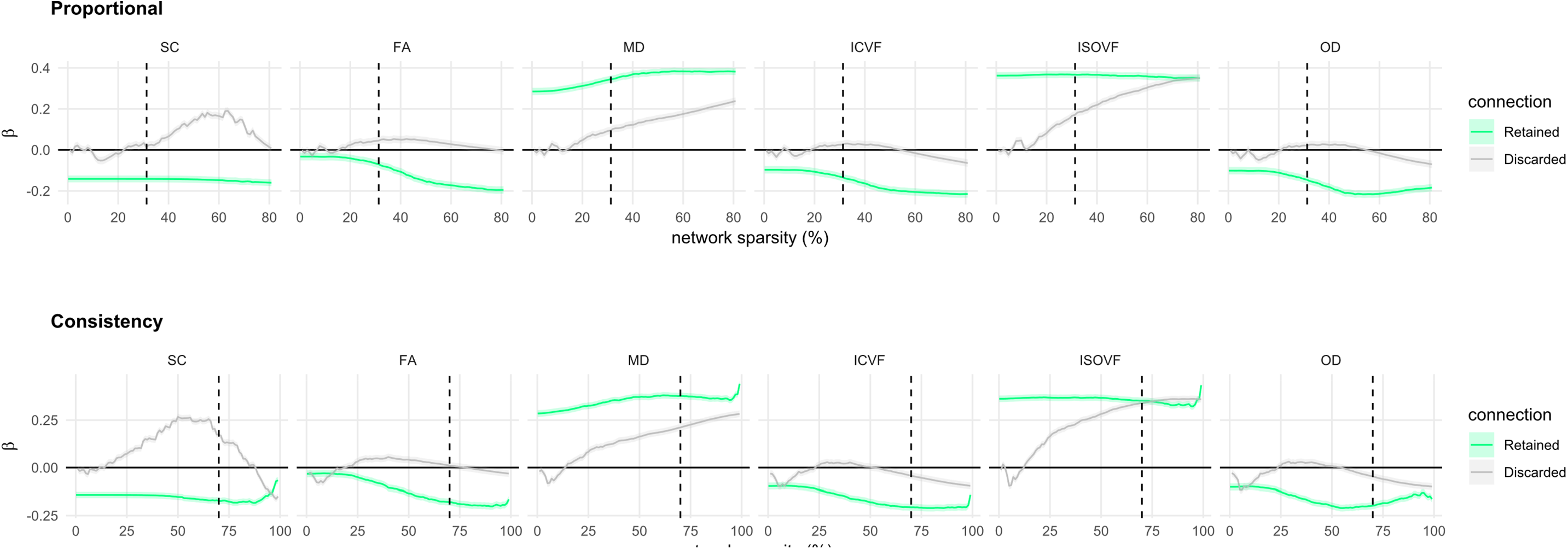
Age-associations (standardised betas with SE) of mean edge weight computed for both connections that were retained and were discarded and over a range of thresholds (0 to 100%) for each of the six network weightings using: proportional-thresholding (top), consistency-thresholding (bottom). The dashed vertical line indicates the threshold levels for 50% of subjects (PT50) and consistency-thresholding at 30% (CT30), respectively.

### Age associations of retained versus discarded connections

Given our hypothesis that connections discarded by thresholding would show mainly null age-associations, we identified the network connections that were both retained and discarded and computed age-associations for each individual connection for both PT50 and CT30 across all six network weightings (Figure 5 and 6). Given that both thresholding methods are agnostic to age, the histograms of age-associations show a marked difference in distribution of retained/discarded connections, particularly for FA, MD, ICVF and ISOVF. Apart from SC weighted networks with PT50 (Figure 5), for all other network weightings and both thresholding approaches the age-associations of the discarded connections (mean β ≤ |0.068|) were significantly smaller in magnitude than the corresponding retained connections (mean β ≤ |0.219|, *p* < 0.001, uncorrected). Interestingly, across all weightings, the distribution of discarded connections (Figure 5A and 6A) had a narrower spread for PT50 (SDs between 0.043 and 0.052) than for the more stringent CT30 method (SDs between 0.065 and 0.085). Examination of the age-associations for each connection (Figures 5B & 6B, bottom row) indicated that CT30 discarded a larger proportion of connections that showed a strong age-association. For CT30, the amount of discarded connections with |β| ≥ 0.20, were 8.2% for ISOVF, 4.4% for MD and ≤ 1.9% for the other weightings.

**Figure 5.**
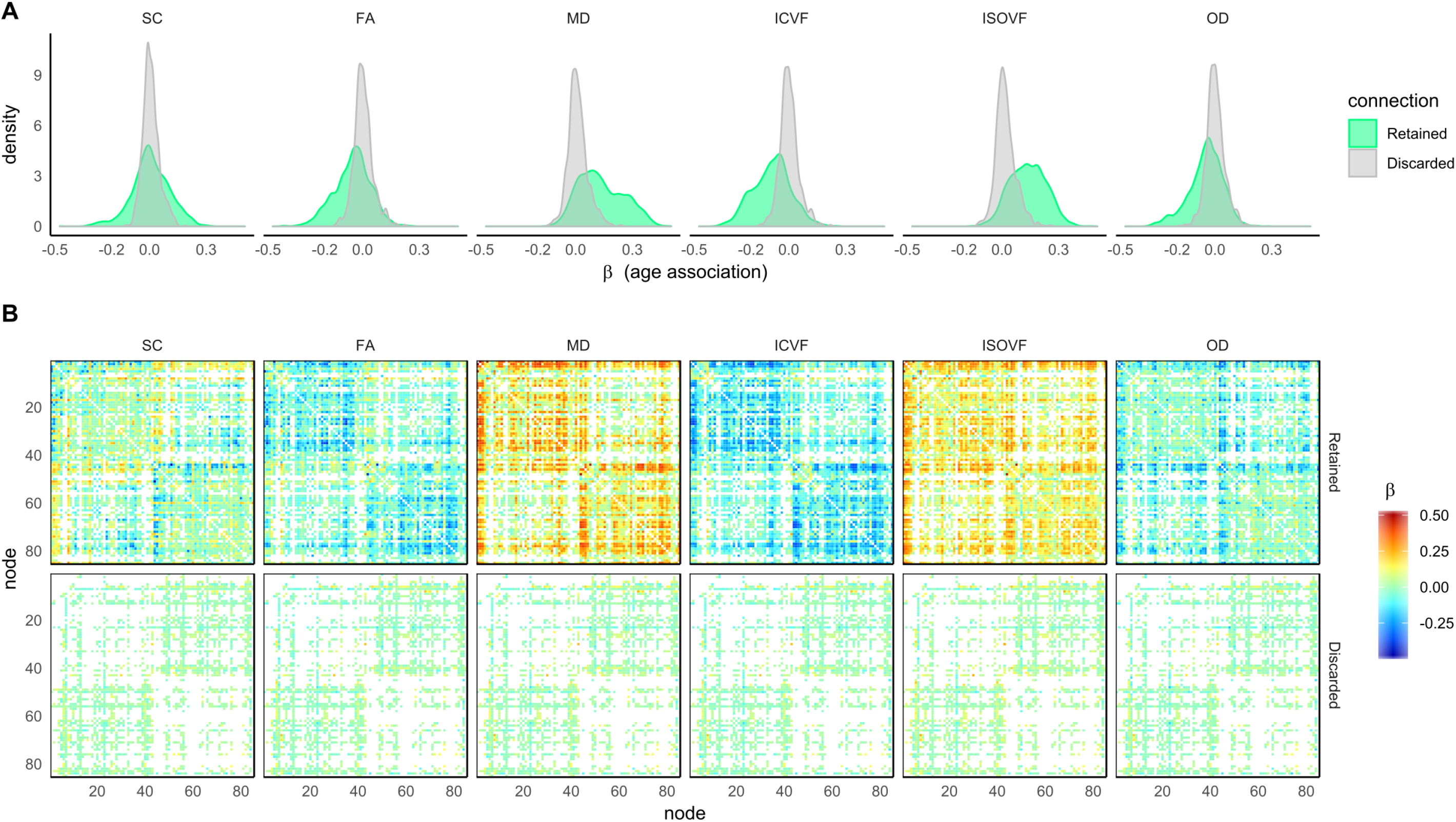
For the six network measures: A) histograms of the age-associations for both connections that were retained and were discarded by proportional-thresholding using 50% of subjects; B) 85 × 85 heatmaps showing the individual age-associations for both connections that were retained and were discarded.

**Figure 6.**
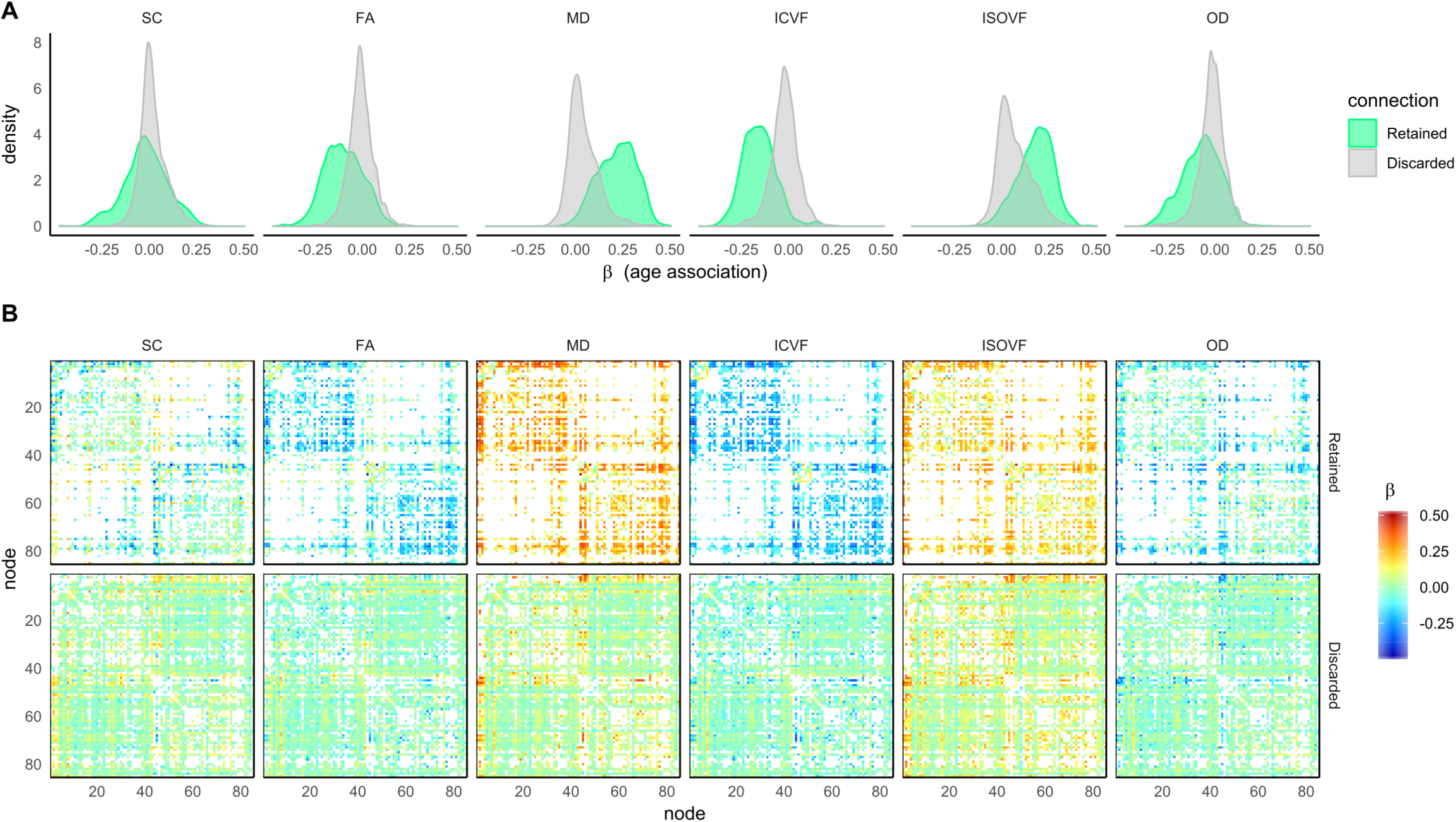
For the six network measures: A) histograms of the age-associations for both connections that were retained and were discarded by consistency-thresholding at 30%; B) 85 × 85 heatmaps showing the individual age-associations for both connections that were retained and were discarded.

In order to identify the regions involved, anatomical circle plots were constructed using CT30 which grouped related neuroanatomical nodes and plotted connections by strength of age-association (*p* < 0.001, uncorrected) for FA and MD weightings (Figure 7; Supplementary Figures 2 and 3 for other weightings). Coherent patterns involving intrahemispheric connections with strong age-associations (|β| ≥ 0.20) were observed for FA, MD and ICVF. For these weightings, the strongest age-associations were for connections between subcortical nodes and connections between frontal nodes. The single strongest age-association was the connection between the left thalamus and left caudate nucleus for MD (β = 0.50). However, SC and to a lesser extend ISOVF and OD showed several interhemispheric connections with strong age-associations. The discarded connections for SC, FA, ICVF and OD (Supplementary Figure 3) involved ≤ 1.9% of connections with relatively strong age-associations (|β| ≥ 0.20). Discarded connections with strong age-associations occurred in somewhat incoherent patterns when compared across weightings and many of the strongest associations involved subcortical connections and connections between subcortical nodes and contralateral cortical nodes.

**Figure 7.**
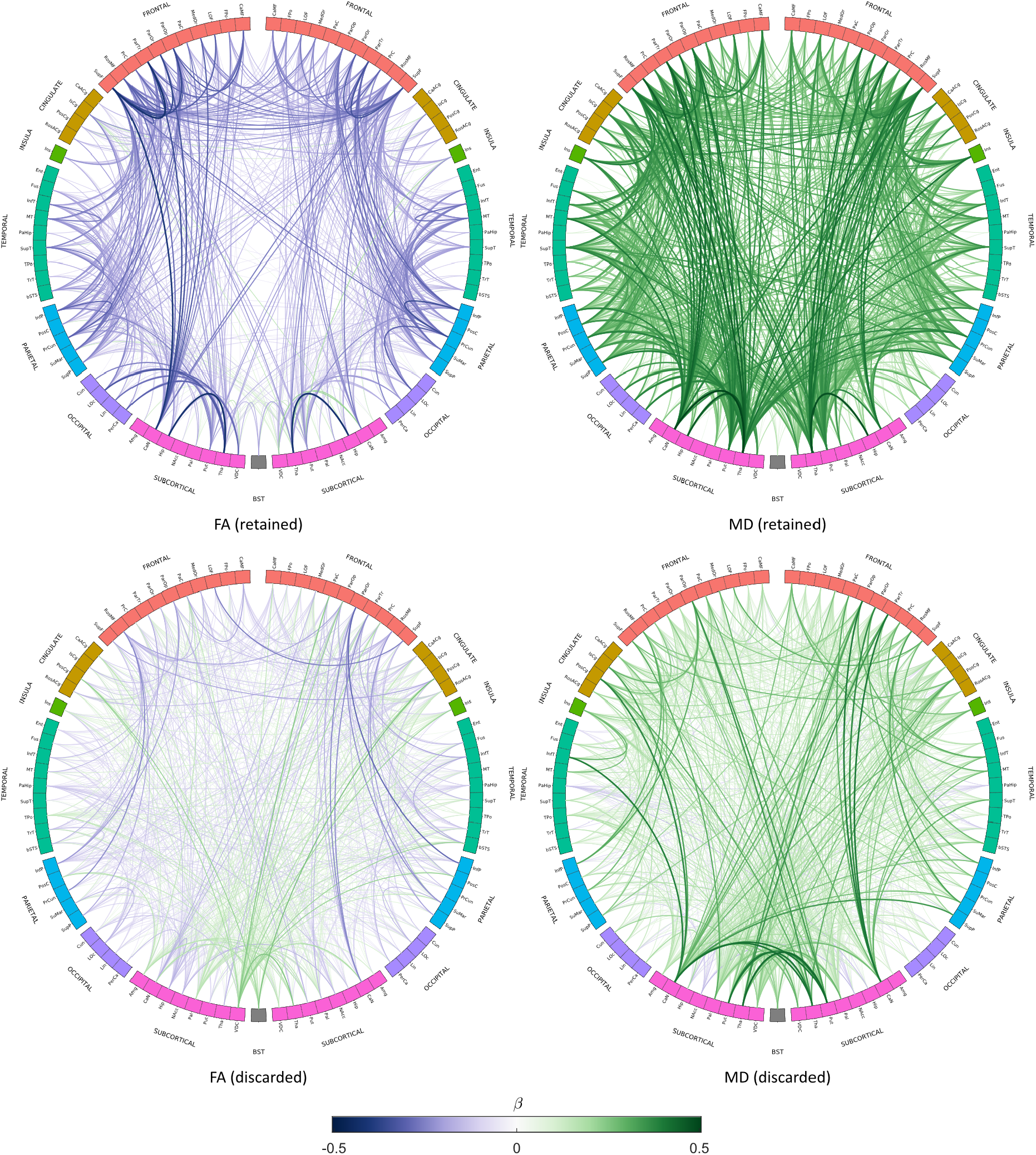
Age-associations (*p* < 0.001, uncorrected) for both retained and discarded connections using consistency-thresholding at 30% for FA- and MD-weighted networks. Link colour represents the age-association (standardised beta) of the mean edge weight, and link thickness represents the magnitude of the association. Node abbreviations are listed in Supplementary Table 7.

## Discussion

This study quantified the effects of principled network thresholding methods and novel network weightings on the criterion validity (associations with age) of an array of structural network metrics. We provided evidence that, in a large sample with previously reported age-associations in white matter (Cox et al., 2016), principled methods to remove implausible connections resulted in the identification of more age-sensitive network components than unthresholded networks. An initial comparison of the group-level thresholding methods (at different levels of sparsity) showed generally stronger age-associations with the more stringent consistency-based (rather than proportional) thresholding. This provides a practical illustration of how currently-used thresholding methods can yield substantially different estimates of association with external variables. However, our more detailed analyses indicated that the stringency of the overall threshold adopted appears to be a stronger determinant of the age-association than the actual method of thresholding itself. When we increased the stringency of the proportional threshold (connections present in 99.6% of individuals, rather than 50%) to match the network sparsity achieved by the consistency-based approach (both at ∼70% network sparsity), both thresholding approaches produced a highly overlapping set of network connections and broadly equivalent age-associations. This finding was echoed when we examined the profile of age-associations across the full sparsity range. There is little difference between the two thresholding approaches in terms of age-associations when sparsity is matched. Generally, more stringent thresholding (increasing sparsity) resulted in a greater magnitude of age-association for retained connections, which were also predominantly closer to previously reported results using major tract-averaged measurements (Cox et al., 2016) than for Raw matrices.

We further compared the profile of age-associations for both retained and discarded connections across all possible network sparsities. In combination with results from multiple approaches indicating pervasive degeneration of the brain’s white matter connections with adult age (Burzynska et al., 2010; Cox et al., 2016; Salat, 2011), our finding that the strongest age-associations fell between 50% and 95% network sparsity, and that the profile of age-associations within the discarded connections were predominantly null across sparsities is consistent with: 1) prior evidence that the human brain is likely to have a connection density of ∼30% (Roberts et al., 2017); 2) that probabilistic tractography fundamentally over-estimates the number of connections (Roberts et al., 2017), and; 3) that these spurious connections add noise to the signal (Jbabdi and Johansen-Berg, 2011). Moreover, our results are inconsistent with previous reports that topological network properties of connectomes are not significantly altered by the removal of weak connections (Civier et al., 2019).

Notably, the profiles of age-associations among the discarded connections appeared distinct when comparing the more general water diffusion parameters (MD and ISOVF) with those metrics thought to convey more specific white matter microstructural information (e.g. FA and ICVF and OD), as shown in Figure 4, for example. The discarded connections showed null age-associations profiles for FA, ICVF and OD across most of the full sparsity range. This is in line with the hypothesis that discarding spurious connections would increase the signal-to-noise ratio. Conversely, the discarded connections weighted by metrics describing the magnitude of general water molecular diffusion (MD and ISOVF) carried significant information about age differences across the majority of thresholds. Diffusion metrics that describe water directional coherence (rather than its general magnitude) should theoretically be more sensitive to the assumed phenomenon of erroneous white matter pathway identification arising from biases in probabilistic tractography. This offers an interesting perspective on differences in the utility of thresholding as a function of the water diffusion metric of interest.

As network sparsity increased, we also found that a larger number of connections with stronger age-associations were discarded. Our investigation of the comparative anatomy of these retained and discarded tracts indicated that the former described coherent bilateral patterns of mainly interhemispheric connectivity, particularly between subcortical nodes and between frontal nodes (FA, MD and ICVF). In addition, age-associations with ipsilateral connections were generally stronger than contralateral connections. In contrast, few of the discarded connections showed strong age-associations, and these were less symmetrical and often involved connections between subcortical and contralateral cortical nodes. Thus, the discarded connections most strongly associated with age were less consistent with findings from mammalian cerebral connectivity (Funnell et al., 2000; Goulas et al., 2017; Oh et al., 2014). Nevertheless, the presence of some contralateral subcortico-cortical connectivity in the mouse (e.g. Oh et al., 2014) could also indicate that more stringent thresholding is increasingly likely to remove some non-spurious connections. Furthermore, some proportion of false-positive connections were likely due to systematic limitations in processing, e.g. an ROI lying close to a major white matter tract which is then wrongly identified as the start/end point of a white matter connection. False continuation or premature termination of streamlines is a known limitation of probabilistic tractography methods (Yeh et al., 2018).

Although various interregional streamline count weightings have previously been used (Buchanan et al., 2014; Hagmann et al., 2008), we have limited our analysis to uncorrected streamlines counts. Our results suggest that streamline-based weightings are affected by volume effects (e.g. the large sex difference with uncorrected SC was likely due to differences in head size and tissue volume). However, streamline count variants that correct for white matter volume and/or grey matter volumes may overcompensate for volume driven effects resulting in age trends in opposite (and unexpected) direction. Some researchers have previously suggested that volume correction of streamline weightings may overcompensate for volume-driven effects on streamline counts (Van Den Heuvel and Sporns, 2011). Therefore, it remains unclear how best to correct streamline weights in order to measure between-subject differences in connectivity. However, the dMRI based weightings, such as FA or MD, are largely agnostic to brain size because they measure the mean value along interconnecting streamlines. No significant sex effects were found with dMRI weightings when measuring mean edge weight.

Beyond our contributions to thresholding and network weighting methods literature, we also contribute robust estimates of associations between age and graph-theoretic metrics which have not previously been reported in a sample of this size. For example, network efficiency for MD and ISOVF-weighted networks were the most age-sensitive measures across a broad range of thresholds; this was considerably stronger than the previously reported latent factor of microstructure from 22 tracts in this sample for these measures (Cox et al., 2016). In addition, the mean edge weight measures of ISOVF and OD were also larger than these previously reported estimates. This could indicate that accounting for a much larger range of white matter connections might provide a clearer reflection of brain-wide white matter correlates of age. Furthermore, we add to our understanding of age differences in white matter microstructure by assessing many more connections than previously reported in this large, single-scanner sample. For example, the previously reported importance of the thalamic radiations for ageing (Cox et al., 2016) could well have been due to sampling error (i.e., bias in the selection of tracts used in the dMRI analysis method did not include other subcortical connections). Whereas our analyses do indicate that thalamic-hippocampal and thalamo-cortical connections exhibit among the strongest age-associations, comparatively strong age-associations were also apparent for caudate and putamen, in line with extant data on the neurostructural underpinnings of certain aspects of cognitive ageing (Fjell et al., 2016).

## Limitations

The study has some limitations. Age-associations are an indirect means against which to verify network properties and should not be used in isolation as a means to detect or prescribe an optimal network thresholding level (e.g., “the largest age-association was found at X% sparsity”), and we do not do so here. Given that thresholding methods are agnostic to age, our use of a descriptive schema allowed us to compare the criterion validity of methods already in use, some of which are substantiated by important biological/histological evidence (Salat, 2011). We also caution that, although age is a well-known correlate of white matter microstructure, there are clearly other plausible explanations by which discarded connections may represent real anatomical pathways that simply do not show age-associations. For comparison, a previous voxel-wise analysis of UK Biobank imaging participants found that, although FA generally decreased with age, some voxels exhibited an increase in FA with age, which may reflect degradation of secondary fibres or reduced fibre dispersion (Miller et al., 2016b). The sample used here comprised generally healthy middle- and older-age participants, who are range restricted in several ways (Fry et al., 2017), which may well affect the generalisability of our findings to other samples. Further methodological work comparing the criterion validity of different network thresholding approaches in the context of pathological neurodegenerative conditions would offer an additional perspective on the current findings.

The value of network variables are known to be sensitive to the construction methodology applied, but there is not yet an agreed schema for constructing structural brain networks from dMRI data (Qi et al., 2015). Necessarily, we have limited our analyses to one approach, but we realise the effect of thresholding will likely differ with other network methods. In particular, principled methods of dMRI denoising, tractography (Tournier et al., 2012) and streamline filtering (Smith et al., 2015, 2013) may further reduce the likelihood of false connections prior to thresholding. Additionally, we selected a brain parcellation schema based on its common application in brain network analyses (Desikan et al., 2006). We conjecture that the biological justification for thresholding at ∼30% connection density (Roberts et al., 2017) will also translate to different neuroanatomical atlases and network resolutions. Nevertheless, the influence of biological and methodological factors remains to be determined across alternative atlases (Qi et al., 2015), particularly those that provide a substantially greater level of granularity (Glasser et al., 2016).

It is challenging to ensure that an arbitrary threshold removes spurious connections while retaining genuine patterns of connectivity. We have limited our analyses to group-level thresholding; consequently, our findings do not necessarily apply to methods that threshold at the individual level, such as absolute-thresholding (Hagmann et al., 2008). Moreover, idiosyncrasies in the structural networks across individuals may limit the utility of such nomothetic approaches. Consistency-thresholding is based on the assumption that connections with the highest inter-subject variability (of corrected streamline counts) are spurious (Roberts et al., 2017). Arguably, removing such connections could hinder the identification of individual differences to be correlated with external variables. Although any network thresholding method is imperfect and can remove genuine patterns of connectivity (at least in a subset of individuals), we believe that thresholding used at plausible levels of network sparsity does remove more false-positive than genuine connections.

## Conclusions

We used a large, single-scanner sample of generally healthy middle- and older-aged participants to test the effects of thresholding and novel network weightings on the criterion validity of network metrics with respect to age. Consistent with biological evidence that the human brain has a connection density of approximately 30%, and the hypothesis that discarded connections would not convey significant information about ageing, we found that more stringent thresholding yielded stronger age-associations than unthresholded networks. This was particularly true for network weightings that denoted directional information (FA, ICVF, OD). Importantly, we found that the specific threshold level applied was a stronger driver of the age-association than the choice of thresholding method.

## Acknowledgments

We thank Clara Alloza and Manuel Blesa Cábez for helpful discussion and insights. We thank the UK Biobank participants and the UK Biobank team for their work in collecting, processing and disseminating these data for analysis. This research was conducted, using the UK Biobank Resource under approved project 10279, in The University of Edinburgh Centre for Cognitive Ageing and Cognitive Epidemiology (CCACE) (http://www.ccace.ed.ac.uk), part of the cross-council Lifelong Health and Wellbeing Initiative (MR/K026992/1). Funding from the Biotechnology and Biological Sciences Research Council (BBSRC) and Medical Research Council (MRC) is gratefully acknowledged. CRB, SRC, SJR, JM, MEB, IJD and EMT-D were supported by a National Institutes of Health (NIH) research grant R01AG054628. SRC and IJD were supported by MRC grants MR/M013111/1 and MR/R024065/1. IJD is additionally supported by the Dementias Platform UK (MR/L015382/1), and he, SRC and SJR receive funding from the Age UK-funded Disconnected Mind project (http://www.disconnectedmind.ed.ac.uk). EMT-D is a member of the Population Research Center at the University of Texas at Austin, which is supported by NIH center grant P2CHD042849.

