## Supplementary Material for "The effect of network thresholding and weighting on structural brain networks in the UK Biobank"

Colin R. Buchanan, Mark E. Bastin, Stuart J. Ritchie, David C. Liewald, James Madole, Elliot M. Tucker-Drob, Ian J. Deary, Simon R. Cox

### Tables

| Title | Notes | Number of participants | Number of nodes | Tractography type | Weighting | Proportional threshold level (%) |
|---|---|---|---|---|---|---|
| (Jung et al., 2017) | Applied a two-step threshold: absolute weight threshold and then proportional threshold at stringent (75%) and relaxed (50%) levels | 24 | 43 | probabilistic | connection probability | 50, 75 |
| (Beare et al., 2017) | The statistical analysis (network-based statistics) used connections that were present in 50% of subjects | 45 | 162 | deterministic, probabilistic | streamline count, weighted streamline count, count fraction | 50 |
| (Nomi et al., 2018) | Applied a proportional threshold at stringent (75%) and relaxed (50%) levels | 199 | 114 | deterministic | streamline count | 50, 75 |
| (Conti et al., 2017) | Applied a two-step threshold: absolute streamline count threshold and connections that were present in all participants | 90 | 48 | probabilistic | streamline count, FA | 100 |
| (Barbagallo et al., 2017) | Applied two alternative thresholds: 1) proportional threshold at 50% ; and 2) an absolute threshold over a range of connection probabilities | 93 | 116 | probabilistic | weighted streamline count | 50 |

**Supplementary Table 1.** Recent network studies using proportional-thresholding were identified by examining the first 100 results since 2017 that matched the search term, "(proportional OR consensus OR group) threshold structural brain network" (search performed using Google Scholar on 13th March 2019) and then identifying those studies that used proportional-thresholding on structural networks constructed from dMRI and tractography (the great majority of search results related to functional networks). For the above studies, the median value of the proportional threshold levels used was a threshold of 50% of subjects.

| Network metric | Weighting | Raw | Proportional (50%) | Consistency (68.6%) | Proportional (99.6%) | Consistency (30%) |
|---|---|---|---|---|---|---|
| | | **sparsity = 0.002** | **sparsity = 0.313** | **sparsity = 0.313** | **sparsity = 0.701** | **sparsity = 0.700** |
| Mean edge weight | SC | 0.005 (0.001) | 0.008 (0.001) | 0.008 (0.001) | 0.017 (0.002) | 0.016 (0.002) |
| | FA | 0.327 (0.020) | 0.435 (0.021) | 0.428 (0.021) | 0.457 (0.016) | 0.460 (0.016) |
| | MD | 0.001 (0.000) | 0.001 (0.000) | 0.001 (0.000) | 0.001 (0.000) | 0.001 (0.000) |
| | ICVF | 0.391 (0.028) | 0.522 (0.032) | 0.514 (0.032) | 0.567 (0.028) | 0.565 (0.028) |
| | ISOVF | 0.058 (0.008) | 0.076 (0.010) | 0.075 (0.010) | 0.078 (0.011) | 0.078 (0.011) |
| | OD | 0.126 (0.007) | 0.169 (0.007) | 0.166 (0.007) | 0.199 (0.007) | 0.196 (0.007) |
| Characteristic path length | SC | 80.029 (14.630) | 80.035 (14.637) | 80.038 (14.638) | 81.079 (17.409) | 80.560 (14.794) |
| | FA | 2.604 (0.111) | 2.717 (0.102) | 2.752 (0.104) | 3.727 (0.123) | 3.682 (0.120) |
| | MD | 1533.703 (57.460) | 1599.896 (54.738) | 1617.745 (55.190) | 2122.544 (74.433) | 2117.128 (71.464) |
| | ICVF | 2.230 (0.126) | 2.329 (0.122) | 2.355 (0.124) | 3.115 (0.150) | 3.093 (0.149) |
| | ISOVF | 12.715 (1.490) | 13.454 (1.655) | 13.747 (1.703) | 21.558 (3.400) | 21.381 (3.194) |
| | OD | 6.597 (0.253) | 6.805 (0.252) | 6.848 (0.255) | 8.626 (0.307) | 8.504 (0.303) |
| Network efficiency | SC | 0.021 (0.003) | 0.021 (0.003) | 0.021 (0.003) | 0.021 (0.003) | 0.021 (0.003) |
| | FA | 0.413 (0.016) | 0.400 (0.015) | 0.397 (0.015) | 0.303 (0.010) | 0.306 (0.010) |
| | MD | 0.001 (0.000) | 0.001 (0.000) | 0.001 (0.000) | 0.001 (0.000) | 0.001 (0.000) |
| | ICVF | 0.487 (0.026) | 0.471 (0.024) | 0.468 (0.024) | 0.364 (0.018) | 0.365 (0.018) |
| | ISOVF | 0.089 (0.010) | 0.085 (0.010) | 0.084 (0.010) | 0.059 (0.008) | 0.059 (0.008) |
| | OD | 0.164 (0.006) | 0.160 (0.006) | 0.159 (0.006) | 0.132 (0.005) | 0.133 (0.005) |
| Network clustering coefficient | SC | 0.002 (0.000) | 0.002 (0.000) | 0.002 (0.000) | 0.007 (0.001) | 0.007 (0.001) |
| | FA | 0.386 (0.016) | 0.377 (0.014) | 0.376 (0.014) | 0.327 (0.011) | 0.309 (0.011) |
| | MD | 0.001 (0.000) | 0.001 (0.000) | 0.001 (0.000) | 0.001 (0.000) | 0.001 (0.000) |
| | ICVF | 0.464 (0.026) | 0.457 (0.024) | 0.455 (0.024) | 0.411 (0.020) | 0.385 (0.019) |
| | ISOVF | 0.063 (0.009) | 0.062 (0.009) | 0.062 (0.009) | 0.050 (0.008) | 0.048 (0.007) |
| | OD | 0.149 (0.006) | 0.149 (0.006) | 0.149 (0.006) | 0.151 (0.005) | 0.140 (0.005) |

**Supplementary Table 2.** Mean (and SD) values of four global network metrics measured across six network weightings and five thresholding approaches. Note that the absolute values of network measures are dependent on network weighting and sparsity. Mean edge weight is the mean of connection weights which survive thresholding. The characteristic path length is a measure of network integration, which reflects the average weighted path length between all pairs of nodes. Network efficiency is an alternative measure of network integration. Network clustering coefficient reflects the average of the local clustering coefficients.

| Network metric | Weighting | Age | Age^2 | Sex | Sex*Age | R^2 |
|---|---|---|---|---|---|---|
| Mean edge weight | SC | **-0.141 (<0.001)** | 0.009 (0.579) | **0.383 (<0.001)** | -0.015 (0.376) | 0.168 |
| | FA | -0.033 (0.079) | -0.001 (0.954) | 0.005 (0.840) | 0.025 (0.181) | 0.000 |
| | MD | **0.285 (<0.001)** | 0.044 (0.009) | 0.055 (0.022) | -0.013 (0.455) | 0.078 |
| | ICVF | **-0.097 (<0.001)** | -0.031 (0.080) | -0.006 (0.809) | 0.019 (0.302) | 0.008 |
| | ISOVF | **0.362 (<0.001)** | **0.068 (<0.001)** | 0.045 (0.054) | 0.005 (0.790) | 0.124 |
| | OD | **-0.101 (<0.001)** | **-0.085 (<0.001)** | 0.005 (0.842) | -0.016 (0.376) | 0.013 |
| Characteristic path length | SC | **0.160 (<0.001)** | 0.019 (0.260) | **-0.231 (<0.001)** | 0.015 (0.410) | 0.077 |
| | FA | **0.089 (<0.001)** | 0.007 (0.699) | -0.030 (0.221) | -0.034 (0.062) | 0.008 |
| | MD | **-0.332 (<0.001)** | **-0.056 (<0.001)** | **-0.089 (<0.001)** | 0.010 (0.553) | 0.109 |
| | ICVF | **0.149 (<0.001)** | 0.043 (0.014) | -0.003 (0.896) | -0.021 (0.239) | 0.020 |
| | ISOVF | **-0.339 (<0.001)** | **-0.064 (<0.001)** | **-0.087 (<0.001)** | -0.023 (0.190) | 0.115 |
| | OD | **0.217 (<0.001)** | **0.115 (<0.001)** | 0.021 (0.388) | 0.024 (0.182) | 0.050 |
| Network efficiency | SC | **-0.186 (<0.001)** | 0.004 (0.792) | **0.381 (<0.001)** | -0.033 (0.048) | 0.182 |
| | FA | **-0.095 (<0.001)** | 0.003 (0.861) | -0.005 (0.847) | 0.042 (0.021) | 0.010 |
| | MD | **0.385 (<0.001)** | **0.077 (<0.001)** | 0.072 (0.002) | -0.012 (0.464) | 0.144 |
| | ICVF | **-0.160 (<0.001)** | -0.037 (0.032) | -0.019 (0.435) | 0.026 (0.149) | 0.024 |
| | ISOVF | **0.380 (<0.001)** | **0.089 (<0.001)** | 0.070 (0.003) | 0.028 (0.105) | 0.142 |
| | OD | **-0.217 (<0.001)** | **-0.118 (<0.001)** | -0.031 (0.197) | -0.020 (0.274) | 0.050 |
| Network clustering coefficient | SC | **-0.098 (<0.001)** | 0.007 (0.686) | **0.385 (<0.001)** | -0.024 (0.162) | 0.160 |
| | FA | -0.056 (0.002) | 0.005 (0.755) | -0.024 (0.343) | 0.036 (0.048) | 0.005 |
| | MD | **0.409 (<0.001)** | **0.073 (<0.001)** | 0.036 (0.119) | -0.019 (0.258) | 0.156 |
| | ICVF | **-0.127 (<0.001)** | -0.034 (0.049) | -0.031 (0.206) | 0.025 (0.175) | 0.016 |
| | ISOVF | **0.345 (<0.001)** | **0.064 (<0.001)** | 0.037 (0.116) | 0.002 (0.930) | 0.111 |
| | OD | **-0.142 (<0.001)** | **-0.109 (<0.001)** | -0.039 (0.109) | -0.023 (0.213) | 0.026 |

**Supplementary Table 3**. Standardised regression coefficients (uncorrected *p*-values) and adjusted $R^2$ measuring mean edge weight, characteristic path length, network efficiency and network clustering coefficient for six network weightings. Computed with unthresholded networks.

| Network metric | Weighting | Age | Age^2 | Sex | Sex*Age | R^2 |
|---|---|---|---|---|---|---|
| Mean edge weight | SC | **-0.141 (<0.001)** | 0.009 (0.580) | **0.383 (<0.001)** | -0.015 (0.375) | 0.168 |
| | FA | **-0.070 (<0.001)** | -0.012 (0.478) | -0.004 (0.859) | 0.028 (0.126) | 0.005 |
| | MD | **0.346 (<0.001)** | 0.045 (0.007) | 0.054 (0.021) | -0.021 (0.224) | 0.113 |
| | ICVF | **-0.134 (<0.001)** | -0.045 (0.009) | -0.018 (0.474) | 0.019 (0.292) | 0.017 |
| | ISOVF | **0.367 (<0.001)** | **0.065 (<0.001)** | 0.047 (0.042) | 0.004 (0.821) | 0.127 |
| | OD | **-0.146 (<0.001)** | **-0.117 (<0.001)** | -0.021 (0.402) | -0.025 (0.177) | 0.028 |
| Characteristic path length | SC | **0.160 (<0.001)** | 0.019 (0.260) | **-0.231 (<0.001)** | 0.015 (0.408) | 0.077 |
| | FA | **0.109 (<0.001)** | 0.010 (0.577) | -0.035 (0.160) | -0.035 (0.055) | 0.012 |
| | MD | **-0.358 (<0.001)** | **-0.057 (<0.001)** | **-0.090 (<0.001)** | 0.009 (0.606) | 0.126 |
| | ICVF | **0.167 (<0.001)** | 0.049 (0.005) | -0.000 (0.992) | -0.022 (0.234) | 0.025 |
| | ISOVF | **-0.341 (<0.001)** | **-0.059 (<0.001)** | **-0.103 (<0.001)** | -0.029 (0.092) | 0.119 |
| | OD | **0.229 (<0.001)** | **0.121 (<0.001)** | 0.033 (0.169) | 0.028 (0.122) | 0.056 |
| Network efficiency | SC | **-0.186 (<0.001)** | 0.004 (0.792) | **0.381 (<0.001)** | -0.033 (0.048) | 0.182 |
| | FA | **-0.117 (<0.001)** | -0.000 (0.985) | -0.004 (0.887) | 0.044 (0.016) | 0.014 |
| | MD | **0.405 (<0.001)** | **0.077 (<0.001)** | 0.072 (0.002) | -0.012 (0.473) | 0.158 |
| | ICVF | **-0.176 (<0.001)** | -0.043 (0.014) | -0.023 (0.359) | 0.026 (0.149) | 0.029 |
| | ISOVF | **0.389 (<0.001)** | **0.087 (<0.001)** | **0.079 (<0.001)** | 0.031 (0.064) | 0.151 |
| | OD | **-0.225 (<0.001)** | **-0.124 (<0.001)** | -0.044 (0.069) | -0.022 (0.210) | 0.055 |
| Network clustering coefficient | SC | **-0.087 (<0.001)** | 0.012 (0.450) | **0.366 (<0.001)** | -0.020 (0.252) | 0.145 |
| | FA | **-0.107 (<0.001)** | -0.000 (0.987) | -0.016 (0.517) | 0.038 (0.039) | 0.013 |
| | MD | **0.406 (<0.001)** | **0.071 (<0.001)** | 0.045 (0.051) | -0.022 (0.197) | 0.155 |
| | ICVF | **-0.162 (<0.001)** | -0.042 (0.016) | -0.027 (0.268) | 0.024 (0.188) | 0.025 |
| | ISOVF | **0.338 (<0.001)** | **0.061 (<0.001)** | 0.046 (0.051) | 0.002 (0.913) | 0.108 |
| | OD | **-0.178 (<0.001)** | **-0.121 (<0.001)** | -0.047 (0.055) | -0.025 (0.170) | 0.039 |

**Supplementary Table 4**. Standardised regression coefficients (uncorrected *p*-values) and adjusted $R^2$ measuring mean edge weight, characteristic path length, network efficiency and network clustering coefficient for six network weightings. Computed with proportional-thresholding using connections present in 50% of subjects.

| Network metric | Weighting | Age | Age^2 | Sex | Sex*Age | R^2 |
|---|---|---|---|---|---|---|
| Mean edge weight | SC | **-0.172 (<0.001)** | 0.000 (0.997) | **0.379 (<0.001)** | -0.022 (0.192) | 0.172 |
| | FA | **-0.179 (<0.001)** | 0.004 (0.800) | -0.012 (0.635) | 0.047 (0.009) | 0.033 |
| | MD | **0.378 (<0.001)** | **0.072 (<0.001)** | 0.048 (0.039) | -0.022 (0.203) | 0.135 |
| | ICVF | **-0.206 (<0.001)** | -0.044 (0.010) | -0.021 (0.400) | 0.026 (0.145) | 0.040 |
| | ISOVF | **0.353 (<0.001)** | **0.062 (<0.001)** | 0.057 (0.015) | 0.003 (0.844) | 0.119 |
| | OD | **-0.198 (<0.001)** | **-0.126 (<0.001)** | -0.034 (0.158) | -0.024 (0.185) | 0.045 |
| Characteristic path length | SC | **0.160 (<0.001)** | 0.020 (0.227) | **-0.230 (<0.001)** | 0.018 (0.312) | 0.076 |
| | FA | **0.140 (<0.001)** | -0.004 (0.799) | -0.051 (0.040) | -0.031 (0.093) | 0.021 |
| | MD | **-0.364 (<0.001)** | **-0.062 (<0.001)** | **-0.081 (<0.001)** | 0.010 (0.570) | 0.129 |
| | ICVF | **0.184 (<0.001)** | 0.043 (0.014) | -0.007 (0.785) | -0.020 (0.282) | 0.031 |
| | ISOVF | **-0.290 (<0.001)** | -0.050 (0.003) | **-0.137 (<0.001)** | -0.022 (0.201) | 0.095 |
| | OD | **0.219 (<0.001)** | **0.121 (<0.001)** | 0.020 (0.409) | 0.026 (0.153) | 0.052 |
| Network efficiency | SC | **-0.192 (<0.001)** | -0.000 (0.992) | **0.378 (<0.001)** | -0.036 (0.033) | 0.182 |
| | FA | **-0.158 (<0.001)** | 0.012 (0.481) | 0.007 (0.765) | 0.048 (0.009) | 0.027 |
| | MD | **0.409 (<0.001)** | **0.084 (<0.001)** | 0.066 (0.004) | -0.010 (0.545) | 0.161 |
| | ICVF | **-0.197 (<0.001)** | -0.040 (0.022) | -0.016 (0.518) | 0.026 (0.149) | 0.036 |
| | ISOVF | **0.401 (<0.001)** | **0.080 (<0.001)** | **0.088 (<0.001)** | 0.024 (0.161) | 0.160 |
| | OD | **-0.215 (<0.001)** | **-0.125 (<0.001)** | -0.033 (0.181) | -0.022 (0.222) | 0.050 |
| Network clustering coefficient | SC | **-0.179 (<0.001)** | -0.011 (0.513) | **0.310 (<0.001)** | -0.036 (0.040) | 0.124 |
| | FA | **-0.192 (<0.001)** | 0.011 (0.540) | -0.008 (0.755) | 0.042 (0.021) | 0.038 |
| | MD | **0.368 (<0.001)** | **0.075 (<0.001)** | 0.046 (0.048) | -0.019 (0.274) | 0.128 |
| | ICVF | **-0.209 (<0.001)** | -0.041 (0.016) | -0.011 (0.658) | 0.025 (0.170) | 0.040 |
| | ISOVF | **0.311 (<0.001)** | 0.054 (0.001) | 0.070 (0.003) | 0.003 (0.856) | 0.094 |
| | OD | **-0.178 (<0.001)** | **-0.122 (<0.001)** | -0.024 (0.326) | -0.017 (0.358) | 0.037 |

**Supplementary Table 5.** Standardised regression coefficients (uncorrected *p*-values) and adjusted $R^2$ measuring mean edge weight, characteristic path length, network efficiency and network clustering coefficient for six network weightings. Computed with consistency-thresholding at 30%.

| Threshold type | Weighting | Age | Age^2 | Sex | Sex*Age | R^2 |
|---|---|---|---|---|---|---|
| Raw | SC-uncorrected | **-0.141 (<0.001)** | 0.009 (0.579) | **0.383 (<0.001)** | -0.015 (0.376) | 0.168 |
| | SC-WM | 0.042 (0.022) | 0.056 (0.001) | **-0.159 (<0.001)** | 0.050 (0.006) | 0.027 |
| | SD-GMV | **0.173 (<0.001)** | 0.020 (0.252) | 0.045 (0.066) | 0.025 (0.167) | 0.033 |
| | SD-GMA | **0.137 (<0.001)** | 0.037 (0.032) | **0.082 (<0.001)** | 0.023 (0.201) | 0.027 |
| Proportional (50%) | SC-uncorrected | **-0.141 (<0.001)** | 0.009 (0.580) | **0.383 (<0.001)** | -0.015 (0.375) | 0.168 |
| | SC-WM | 0.042 (0.022) | 0.056 (0.001) | **-0.159 (<0.001)** | 0.050 (0.006) | 0.027 |
| | SD-GMV | **0.173 (<0.001)** | 0.020 (0.253) | 0.045 (0.067) | 0.025 (0.167) | 0.033 |
| | SD-GMA | **0.137 (<0.001)** | 0.037 (0.032) | **0.082 (<0.001)** | 0.023 (0.201) | 0.027 |
| Consistency (30%) | SC-uncorrected | **-0.172 (<0.001)** | 0.000 (0.997) | **0.379 (<0.001)** | -0.022 (0.192) | 0.172 |
| | SC-WM | -0.005 (0.783) | 0.042 (0.015) | **-0.173 (<0.001)** | 0.042 (0.020) | 0.030 |
| | SD-GMV | **0.116 (<0.001)** | -0.004 (0.823) | 0.025 (0.305) | 0.011 (0.531) | 0.014 |
| | SD-GMA | **0.067 (<0.001)** | 0.014 (0.427) | 0.061 (0.015) | 0.007 (0.691) | 0.007 |
| Proportional (99.6%) | SC-uncorrected | **-0.154 (<0.001)** | 0.007 (0.640) | **0.384 (<0.001)** | -0.015 (0.358) | 0.172 |
| | SC-WM | 0.023 (0.217) | 0.054 (0.002) | **-0.160 (<0.001)** | 0.050 (0.006) | 0.027 |
| | SD-GM | **0.146 (<0.001)** | 0.014 (0.430) | 0.036 (0.145) | 0.025 (0.170) | 0.023 |
| | SD-GMA | **0.105 (<0.001)** | 0.032 (0.067) | 0.074 (0.003) | 0.022 (0.226) | 0.017 |
| Consistency (68.6%) | SC-uncorrected | **-0.142 (<0.001)** | 0.009 (0.587) | **0.383 (<0.001)** | -0.015 (0.372) | 0.168 |
| | SC-WM | 0.041 (0.025) | 0.056 (0.001) | **-0.159 (<0.001)** | 0.049 (0.007) | 0.027 |
| | SD-GMV | **0.172 (<0.001)** | 0.019 (0.267) | 0.044 (0.076) | 0.025 (0.170) | 0.032 |
| | SD-GMA | **0.136 (<0.001)** | 0.037 (0.035) | 0.080 (0.001) | 0.023 (0.204) | 0.026 |

**Supplementary Table 6**. Standardised regression coefficients (uncorrected *p*-values) and adjusted $R^2$ measuring mean edge weight for four streamline count network weightings: SC) Uncorrected streamline count; SC-WM) network-wise correction by number of seed points per subject (count of white matter voxels); SD-GM) streamline density with edge-wise correction by node volumes (count of voxels in node ROI); and SD-GMA) streamline density with edge-wise correction by node surface area at the white matter interface (the count of voxels which directly neighbour a white matter voxel).

| Region | Abbreviation | Lobe/Grouping |
|---|---|---|
| Caudal middle frontal | CaMF | Frontal |
| Frontal pole | FPo | Frontal |
| Lateral orbitofrontal | LOF | Frontal |
| Medial orbitofrontal | MedOr | Frontal |
| Paracentral | PaC | Frontal |
| Pars opercularis | ParOp | Frontal |
| Pars orbitalis | ParOr | Frontal |
| Pars triangularis | ParTr | Frontal |
| Precentral | PrC | Frontal |
| Rostral middle frontal | RosMF | Frontal |
| Superior frontal | SupF | Frontal |
| Caudal anterior cingulate | CaACg | Cingulate |
| Isthmus cingulate | IsCg | Cingulate |
| Posterior cingulate | PosCg | Cingulate |
| Rostral anterior cingulate | RosACg | Cingulate |
| Insula | Ins | - |
| Banks of superior temporal | bSTS | Temporal |
| Entorhinal | Ent | Temporal |
| Fusiform | Fus | Temporal |
| Inferior temporal | InfT | Temporal |
| Middle temporal | MT | Temporal |
| Parahippocampal | PaHip | Temporal |
| Superior temporal | SupT | Temporal |
| Temporal pole | TPo | Temporal |
| Transverse temporal | TrT | Temporal |
| Inferior parietal | InfP | Parietal |
| Postcentral | PosC | Parietal |
| Precuneus | PrCun | Parietal |
| Superior parietal | SupP | Parietal |
| Supramarginal | SuMar | Parietal |
| Cuneus | Cun | Occipital |
| Lateral occipital | LOc | Occipital |
| Lingual | Lin | Occipital |
| Pericalcarine | PerCa | Occipital |
| Nucleus accumbens | NAcc | Subcortical |
| Amygdala | Amg | Subcortical |
| Caudate nucleus | CaN | Subcortical |
| Hippocampus | Hip | Subcortical |
| Pallidum | Pal | Subcortical |
| Putamen | Put | Subcortical |
| Thalamus | Tha | Subcortical |
| Ventral diencephalon | VDC | Subcortical |
| Brainstem | BSt | - |

**Supplementary Table 7**. List of neuroanatomical regions and abbreviations.

### Figures

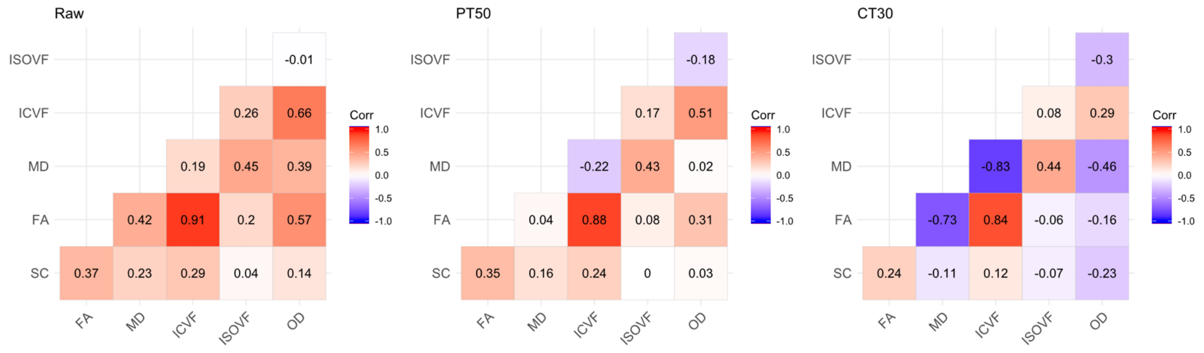


**Supplementary Figure 1.** Heatmap showing correlations between the mean edge weight of the six network weightings for: left) unthresholded matrices; middle) proportional-thresholding at 50% (PT50) of subjects; and consistency-thresholding at 30% (CT30).

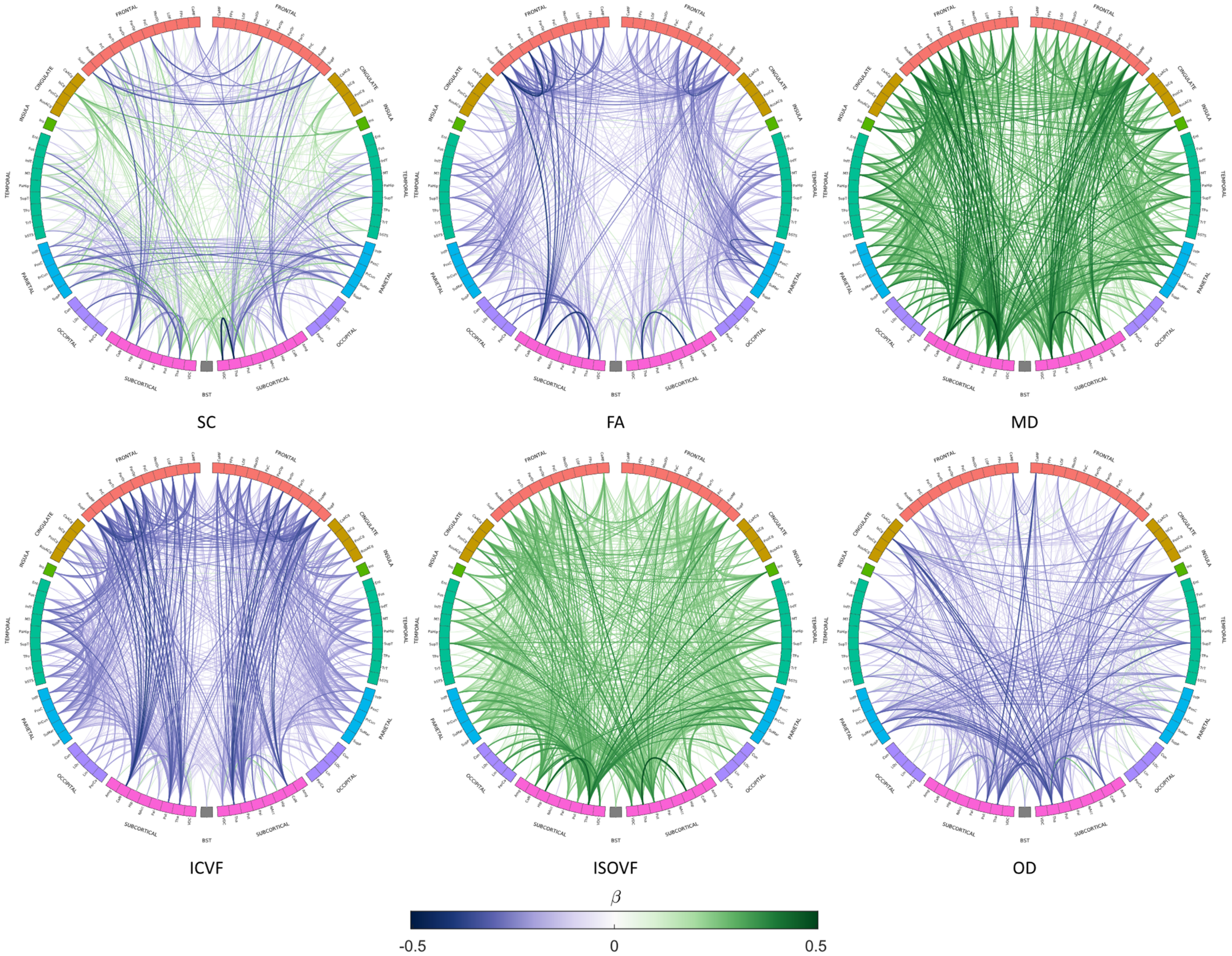


**Supplementary Figure 2**. Age-associations ($p < 0.001$, uncorrected) for connections which survive consistency-thresholding at 30% for 6 network weightings (SC, FA, MD, ICVF, ISOVF and OD). Link colour and thickness represents the age-association (standardised beta) of the mean edge weight.

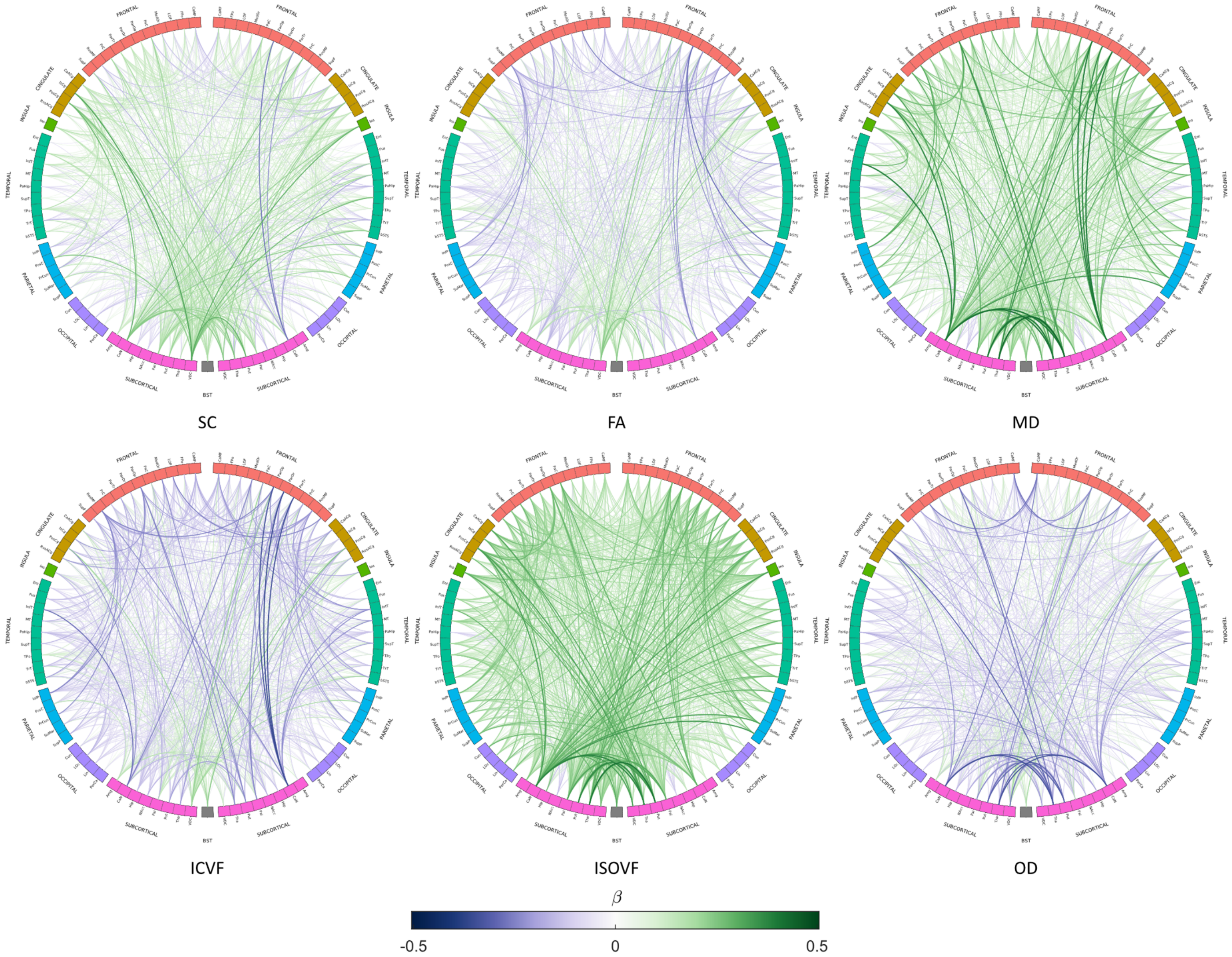


**Supplementary Figure 3.** Age-associations ($p < 0.001$, uncorrected) for connections which are discarded by consistency-thresholding at 30% for 6 network weightings (SC, FA, MD, ICVF, ISOVF and OD). Link colour and thickness represents the age-association (standardised beta) of the mean edge weight.
